# A genetically-defined population in the lateral and ventrolateral periaqueductal gray selectively promotes flight to safety

**DOI:** 10.1101/2022.01.19.476981

**Authors:** Mimi La-Vu, Ekayana Sethi, Sandra Maesta-Pereira, Peter J Schuette, Brooke C Tobias, Fernando MCV Reis, Weisheng Wang, Saskia J Leonard, Lilly Lin, Avishek Adhikari

## Abstract

When encountering external threats, survival depends on the engagement of appropriate defensive reactions to minimize harm. There are major clinical implications for identifying the neural circuitry and activation patterns that produce such defensive reactions, as maladaptive overactivation of these circuits underlies pathological human anxiety and fear responses. A compelling body of work has linked activation of large glutamatergic neuronal populations in the midbrain periaqueductal gray (PAG) to defensive reactions such as freezing, flight and threat-induced analgesia. These pioneering data have firmly established that the overarching functional organization axis of the PAG is along anatomically-defined columnar boundaries. Accordingly, broad activation of the dorsolateral column induces flight, while activation of the lateral or ventrolateral (l and vl) columns induces freezing. However, the PAG contains a diverse arrangement of cell types that vary in neurochemical profile and location. How these cell types contribute to defensive responses remains largely unknown, indicating that targeting sparse, genetically-defined populations can lead to a deeper understanding of how the PAG generates a wide array of behaviors. Though several prior works showed that broad excitation of the lPAG or vlPAG causes freezing, we found that activation of lateral and ventrolateral PAG (l/vlPAG) cholecystokinin-expressing (cck) cells selectively causes flight to safer regions within an environment. Furthermore, inhibition of l/vlPAG-cck cells reduces avoidance of a predatory threat without altering other defensive behaviors like freezing. Lastly, l/vlPAG-cck activity increases away from threat and during movements towards safer locations. In contrast, activating l/vlPAG cells pan-neuronally promoted freezing and these cells were activated near threat. These data underscore the importance of investigating genetically-identified PAG cells. Using this approach, we found a sparse population of cck-expressing l/vlPAG cells that have distinct and opposing function and neural activation motifs compared to the broader local ensemble defined solely by columnar anatomical boundaries. Thus, in addition to the anatomical columnar architecture of the APG, the molecular identity of PAG cells may confer an additional axis of functional organization, revealing unexplored functional heterogeneity.

## Introduction

The midbrain periaqueductal gray (PAG) has been implicated in numerous functions including pain modulation, vocalization, breathing, heart rate, hunting, freezing, and flight (Behbehani, 1995; Keay and Bandler, 2015; Motta et al., 2017; Silva and McNaughton, 2019). For decades, a great deal of effort has been put toward understanding how columnar subdivisions of the PAG control or contribute to distinct defensive behaviors (Bandler et al., 1985; Bandler and Carrive, 1988; Bandler and Shipley, 1994; Carrive, 1993; de Andrade Rufino et al., 2019; Gross and Canteras, 2012; Leman et al., 2003; Morgan and Clayton, 2005; Tomaz et al., 1988; Walker and Carrive, 2003; Zhang et al., 1990). Prior work indicates that the ventrolateral PAG column is necessary for conditioned freezing (Tovote et al., 2016). Though less studied than the vlPAG, optogenetic and electrical excitation of the lateral (l) PAG column also produces freezing (Assareh et al., 2016; Bittencourt et al., 2005; Yu et al., 2021). Moreover, the dorsolateral (dl) PAG has a key role in controlling innate defensive behaviors. Indeed, dlPAG cells encode numerous defense behaviors including freezing, escape and risk-assessment (Del-Ben and Graeff, 2009; Deng et al., 2016; Reis et al., 2021), and activation of glutamatergic vGlut2+ dlPAG cells induces escape (Evans et al., 2018; Tovote et al., 2016). More recent work employing methods with genetic specificity have focused on large PAG populations positive for broadly expressed markers such as *Vgat, Vglut2* or *CaMKIIa*. For example, optogenetic activation of PAG neurons expressing *CaMKIIa*, which is ubiquitous in the region, elicited both freezing and flight. DlPAG cells were also active during escape, threat proximity and risk-assessment (Deng et al., 2016). Though these interesting data provided important insights, they leave open the question of whether sparser PAG populations might control and encode more specific behavioral metrics.

Indeed, the PAG contains a diverse array of sparse cell types (Keay and Bandler, 2015; Silva and McNaughton, 2019; Yin et al., 2014). These cell types exhibit different neurochemical profiles and vary in anatomical location, often spanning more than a single column (Silva and McNaughton, 2019). For example, substance P-producing *Tac1*+ cells and enkephalin-releasing *Penk*+ cells are concentrated in dorsomedial and ventrolateral posterior PAG, while somatostatin-expressing cells can be found in dorsomedial and lateral columns (Allen Brain Atlas 2021; (Silva and McNaughton, 2019). It is possible that distinct cell types contribute to specific phenotypes controlled by the PAG. Accordingly, genetically-identified populations have been more deeply studied in other regions such as the lateral hypothalamus (Li et al., 2018) or the central amygdala (Fadok et al., 2017), leading to unprecedented insights on their function. However, cell-type specific dissections of sparse PAG populations remain scarce, and the functions of specific molecularly-defined cell populations are largely uncharacterized.

One population of interest is composed of cholecystokinin-releasing PAG (PAG-cck) cells (Allen Brain Atlas 2021). Intriguingly, intra-PAG infusion of cck in rats induces defensive behaviors and potentiates one-way escape behavior (Netto and Guimarães, 2004; Zanoveli et al., 2004). Additionally, cck excites PAG neurons at both pre- and postsynaptic loci (Liu et al., 1994; Yang et al., 2006), suggesting PAG-cck cells may have widespread effects on local cell activity dynamics. However, despite these tantalizing results, to date PAG-cck cells have not been directly studied and their function remains unknown.

Here, we specifically target, manipulate and monitor the neural activity of PAG-cck cells. We show that lateral and ventrolateral (l/vl) PAG-cck cells are a small subset of glutamatergic cells, and that they selectively control flight to a safe location within an environment without affecting other defensive behaviors such as freezing or other l/vlPAG-mediated processes such as analgesia. Furthermore, though decades of prior work has consistently shown that PAG cells are activated by proximity to danger (Aguiar and Guimarães, 2009; Canteras and Goto, 1999; Deng et al., 2016; Evans et al., 2018; Mobbs et al., 2010, 2007; Watson et al., 2016), we find that l/vlPAG-cck cells are activated far from threat. In contrast, broad pan-neuronal activation of cells in the same l/vlPAG region induced freezing, and these cells encoded a wider variety of behaviors. Thus, we show that characterization of sparser, genetically-defined PAG populations may reveal cells that have unique functional roles and that may even show opposing patterns of neural activation relative to the broader local ensemble. Deciphering how molecularly-defined PAG populations complement and interact with the well-established anatomical columnar functional framework is a key step in understanding how this ancient structure controls a constellation of vital behaviors.

## Results

### Cck+ cells comprise a sparse glutamatergic subset of l/vlPAG neurons

Cck is expressed primarily in two clusters within the PAG: one located in the dorsomedial column and one spanning both the lateral and ventrolateral columns (Allen Brain Atlas 2021). Here, we focused on the latter population, which is more prevalent in the posterior than anterior PAG. To quantify the proportion of cck+ neurons in the posterior l/vlPAG column, we used cre-dependent viral vectors to express GFP in cck cells of *Cck-ires-cre* mice. We then immunostained posterior PAG slices against the pan-neuronal marker NeuN and quantified Neun/GFP overlap (Figure 1A). We observed GFP expression in the lPAG and vlPAG, but GFP expression was largely absent in the dlPAG (Figure 1B). Quantification showed that cck-GFP cells comprise approximately 5% of l/vlPAG neurons (Figure 1C). Cholecystokinin-expressing cells in several brain regions such as cortex, hippocampus and amygdala are reported to be inhibitory (Kepecs and Fishell, 2014; Mascagni and McDonald, 2003; Nguyen et al., 2020; Whissell et al., 2015), though glutamatergic cck cells have also been reported in other regions (Wang et al., 2021). To determine if these cells are glutamatergic, we immunostained against the glutamatergic marker vGlut2 in PAG slices containing GFP-expressing cck+ cells (Figure 1D). Notably, we found 95% of GFP-labeled cells were also vGlut2-labeled (Figure 1E). Our characterization shows that cck+ cells comprise a small, sparse subset of PAG neurons that span the lateral and the ventrolateral columns and are primarily glutamatergic.

**Figure 1.**
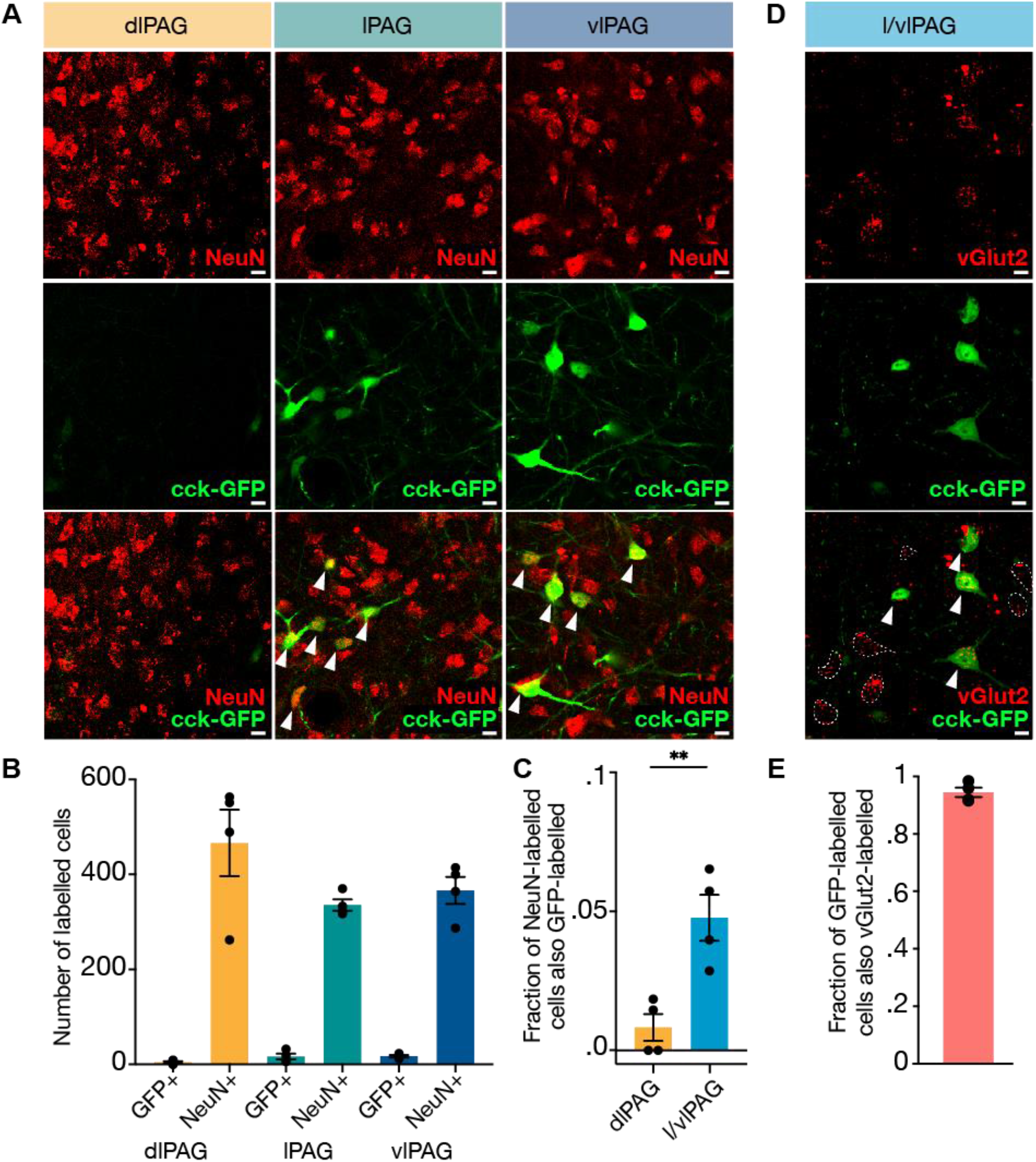
Cck+ cells comprise approximately 5% of l/vlPAG neurons and are primarily glutamatergic. **(A)** Example histology images showing immunostaining of pan-neuronal marker NeuN (top row), viral-mediated expression of GFP in cck-expressing cells (middle row), and overlay of NeuN and cck-GFP (bottom row) in the dorsolateral (left column), lateral (middle column), and ventrolateral (right column) PAG. Scale bars, 10μm. **(B)** Raw counts of cck-GFP+ and NeuN+ cells in the dlPAG, lPAG, and vlPAG. **(C)** Fraction of NeuN-labeled cells that are also GFP-labeled in the dlPAG and l/vlPAG. Cck-expressing cells comprise ∼5% of l/vlPAG neurons and constitute significantly more of l/vlPAG neurons than dlPAG neurons (n=4; paired t-test, **p < .01). **(D)** Immunostaining of glutamatergic marker vGlut2 in cck cells. Example histology images showing vGlut2 (top), cck-GFP (middle) and vGlut2/GFP overlay (bottom). White arrow indicates vGlut2+/GFP+ cell. Dashed outline indicates vGlut2+/GFP-cell. Scale bar, 10μm. **(E)** Approximately 95% of GFP-labeled cells are also vGlut2-labeled (n = 4). Mean ± SEM.

### l/vlPAG-cck stimulation induces a repertoire of behaviors distinct from pan-neuronal dlPAG and l/vlPAG stimulation

To study how various PAG populations may participate in distinct defensive phenotypes, we used an optogenetic approach to manipulate three different PAG subpopulations: pan-neuronal synapsin (syn)-expressing dorsolateral PAG neurons (dlPAG-synapsin), pan-neuronal syn-expressing lateral/ventrolateral PAG neurons (l/vlPAG-synapsin) and cholecystokinin-expressing lateral/ventrolateral PAG neurons (l/vlPAG-cck). We targeted these populations by local injection of adeno-associated viruses (AAVs) delivering channelrhodopsin-2 (ChR2) coupled to a yellow fluorescent protein (YFP) tag into the dlPAG or l/vlPAG of wildtype (WT) mice and l/vlPAG of *Cck-ires-cre mice* (Figure 2A-B). Mice injected with AAVs containing only YFP served as controls. The viral strategy used to transfect pan-neuronal l/vlPAG cells was synapsin-specific and did not exclude transfection of cck+ cells. We first optogenetically manipulated naive mice in an open field (Figure 2C-G). Optogenetic activation of syn-expressing dlPAG cells increased speed and open field corner entries (Figure 2H, L). Notably, activation of only this, but not other PAG populations, induced escape jumping (Figure 2I). Light activation of syn-expressing l/vlPAG cells strongly promoted freezing, and consequently reduced speed and corner entries (Figure 2H, J, L). Finally, we observed that activation of l/vlPAG-cck cells increased speed, reduced time spent in the open field center, and increased corner entries (Figure 2H, K-L). Interestingly, activation of only this population increased time spent in the corners of the open field (Figure 2M). These results demonstrate that increased activity in these three PAG subpopulations elicited diverse behavioral phenotypes. Stimulation of l/vlPAG-cck cells induced a repertoire of behaviors distinct from pan-neuronal l/vlPAG and dlPAG activation. Specifically, activation induced a preference for the corners of the open field, which represent the safest area in the arena as they allow mice to best limit visual detection by predators (La-Vu et al., 2020).

**Figure 2.**
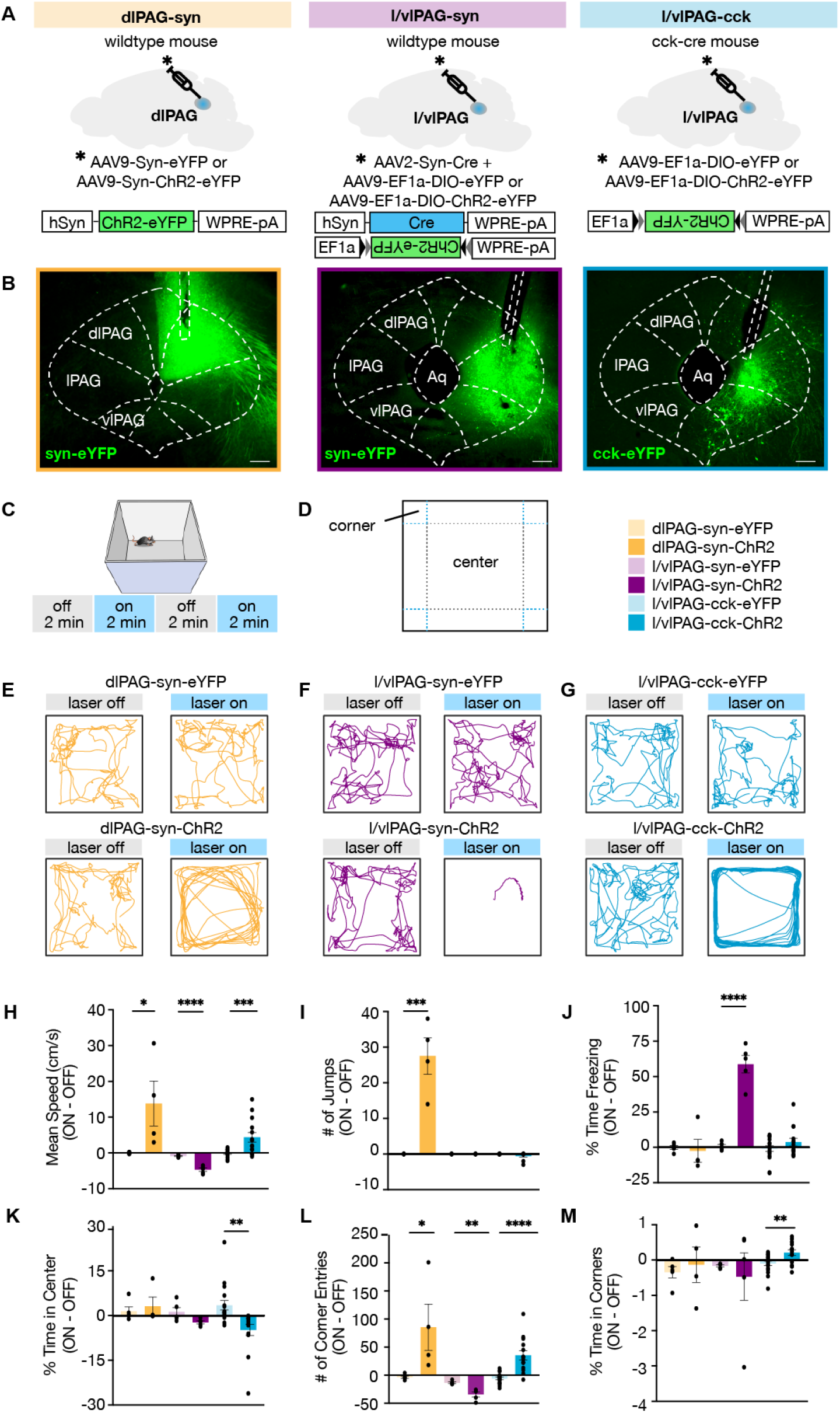
l/vlPAG-cck stimulation induced a repertoire of behaviors distinct from pan-neuronal l/vlPAG and dlPAG activation. **(A)** Viral strategy to express enhanced yellow fluorescent protein (eYFP) or light-sensitive channelrhodopsin (ChR2-eYFP) in synapsin-expressing cells in the dorsolateral periaqueductal grey (dlPAG-syn, left), synapsin-expressing cells in the lateral and ventrolateral PAG (l/vlPAG-syn, middle), and cholecystokinin-expressing cells in the lateral and ventrolateral PAG (l/vlPAG-cck, right). A fiber optic cannula was then implanted over respective regions. **(B)** Histology of eYFP expression in dlPAG-syn (left), l/vlPAG-syn (middle) and l/vlPAG-cck (right). Scale bar, 200μm. **(C)** Stimulation protocol in the Open Field. Blue light (473-nm, 5-ms, 20-Hz) was delivered in alternating 2-min epochs (OFF-ON-OFF-ON) for 8-min total. **(D)** Diagram indicating center and corners of the Open Field assay. **(E-G)** Example locomotion maps in the Open Field during laser-off and laser-on epochs of either mice expressing eYFP (top) or ChR2-eYFP (bottom) in dlPAG-syn (**E**), l/vlPAG-syn (F), and l/vlPAG-cck (**G**) populations. **(H-M)** Bars depict respective behaviors during light-off epochs subtracted from light-on epochs (ON minus OFF). Light delivery to dlPAG of syn-ChR2 mice increased mean speed, jumps and corner entries compared to dlPAG-syn-eYFP mice. Light delivery to the l/vlPAG of syn-ChR2 mice reduced mean speed, increased freezing, and reduced corner entries compared to l/vlPAG-syn-eYFP mice. Light delivery to the l/vlPAG of cck-ChR2 mice increased mean speed and corner entries while reducing center time compared to l/vlPAG-cck-eYFP mice. Importantly, time spent in corners increased during light delivery to l/vlPAG-cck-ChR2 mice compared to l/vlPAG-cck-eYFP mice (dlPAG: eYFP, n = 5, ChR2, n = 4; l/vlPAG-syn: eYFP, n = 5; ChR2, n = 5; l/vlPAG-cck: eYFP, n = 17, ChR2, n = 14; unpaired t-test, *p < 0.05, **p < 0.01, ***p < 0.001, ****p < 0.0001). Mean ± SEM.

### l/vlPAG-cck stimulation prompts entry into a dark burrow

We aimed to further investigate the exhibited preference for safety upon activation of l/vlPAG-cck cells. We developed the Latency to Enter (LTE) assay, a novel paradigm that measures flight to the safest region within an environment. The LTE is a square arena illuminated to 80 lux and contains a dark burrow (2 lux) in one corner. Mice were habituated to the arena for 10 min; only mice that exhibited a preference for the burrow during habituation continued to Test on the following day. During Test, mice were placed in the LTE for a 1-min context reminder prior to ten consecutive trials. For optogenetic manipulation within the LTE, light delivery was alternated across the ten trials, beginning with a light-off trial. Prior to the start of each trial, mice were confined to the corner opposite of the burrow, the holding zone, with a transparent barrier for 15 sec. For light-on trials, light was delivered for the last 5 sec of the 15-sec period in the holding zone and continued until the end of the trial. The start of a trial was marked by barrier removal and the trial ended upon burrow entry or 60 sec had passed. If a mouse entered the burrow, they could remain in the burrow for 10 sec prior to being returned to the holding zone. If they did not enter, they were immediately returned to the holding zone.

To study if activation of PAG subpopulations can bias mice to flee to the burrow, we optogenetically manipulated dlPAG, l/vlPAG, and l/vlPAG-cck neurons in the Latency to Enter assay (Figure 3A-B). Despite similar levels of preference for the burrow during habituation, only optogenetic activation of l/vlPAG-cck neurons reduced latency to enter the burrow relative to YFP control mice (Figure 3C-F). Notably, activation of syn-expressing l/vlPAG neurons robustly increased latency, as l/vlPAG-syn-ChR2 mice displayed substantial freezing with light-delivery compared to YFP mice. These data demonstrate increased activity in l/vlPAG-cck neurons can induce urgent flight to relative safety in a low-threat environment, a feature distinct from pan-neuronal l/vlPAG and dlPAG cells.

**Figure 3.**
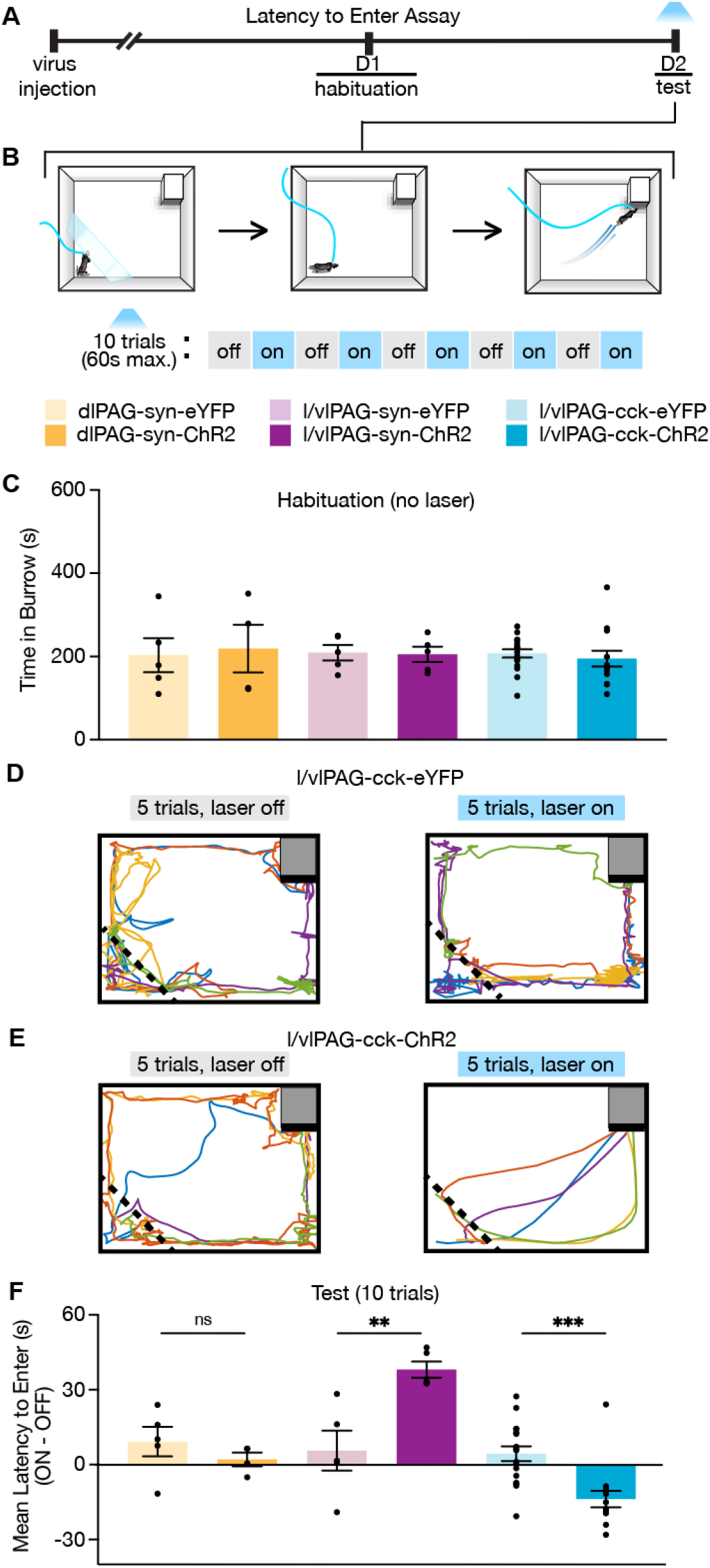
l/vlPAG-cck stimulation prompts entry into a dark burrow in the absence of threat, unlike pan-neuronal l/vlPAG and dlPAG stimulation. **(A)** Timeline of Latency to Enter (LTE) assay. Mice were habituated to a chamber containing a dark burrow for 10-min one day prior to test. Mice with preference for burrow during habituation were included in the test. **(B)** Schematic of LTE assay during Test. Left: At the beginning of each trial, mice were confined to a corner opposite of a dark burrow with a transparent barrier (holding zone). After 15 s, the barrier was removed and mice could freely move about the chamber. The trial ended when mice entered the burrow or 60 s had passed. Blue light (473-nm, 5-ms, 20-Hz) was delivered in alternating trials. In light-on trials, blue light delivery began 5-s prior to barrier removal. After burrow entry, mice could remain in the burrow for 10 s before being returned to the holding zone. **(C)** Bars represent average time spent in the burrow during a 10-min habituation (dlPAG: eYFP, n = 5; ChR2, n = 4; l/vlPAG-syn: eYFP, n = 5; ChR2, n = 5; l/vlPAG-cck: eYFP, n = 17, ChR2, n = 14). **(D-E)** Example locomotion map of 5 light-off (left) and 5 light-on trials (right) in a l/vlPAG-cck-eYFP mouse (**D**) and l/vlPAG-cck-ChR2 mouse (**E**). **(F)** Individual dots represent mean latency during 5 light-on epochs subtracted by mean latency during 5 light-off epochs (ON minus OFF). Trials without entry were regarded as latency of 61 s. Light delivery increased latency to enter the burrow in l/vlPAG-syn-ChR2 mice compared to l/vlPAG-syn-eYFP mice and reduced latency to enter the burrow in l/vlPAG-cck-ChR2 mice compared to l/vlPAG-cck-YFP mice (dlPAG: eYFP, n = 5; ChR2, n = 4; l/vlPAG-syn: eYFP, n = 5; ChR2, n = 5; l/vlPAG-cck: eYFP, n = 17, ChR2, n = 14; unpaired t-test, **p < 0.01, ***p < 0.001). Mean ± SEM.

### l/vlPAG-cck stimulation is aversive and anxiogenic, and induces a hallmark sympathetic response

As there are no reports of genetically-targeted manipulation of l/vlPAG-cck cells, we sought to further characterize the behavioral phenotype induced by l/vlPAG-cck activation. We assessed the effects of optogenetic activation in mice expressing ChR2 in l/vlPAG-cck cells compared to YFP controls in anxiety and defense-related assays. Pairing light activation of l/vlPAG-cck cells to one chamber in the real-time place test assay resulted in avoidance of the stimulated chamber, suggesting increased l/vlPAG-cck activity is aversive (Figure 4A-C). Furthermore, stimulation of l/vlPAG-cck cells in the elevated plus maze (EPM) reduced time spent in the open arms of the maze (Figure 4D-F). Light activation of l/vlPAG-cck cells also markedly increased pupil size, a hallmark sympathetic response (Figure 4G-H). Together, these results suggest l/vlPAG-cck cell activation is aversive, anxiogenic, and elicits sympathetic activation.

**Figure 4.**
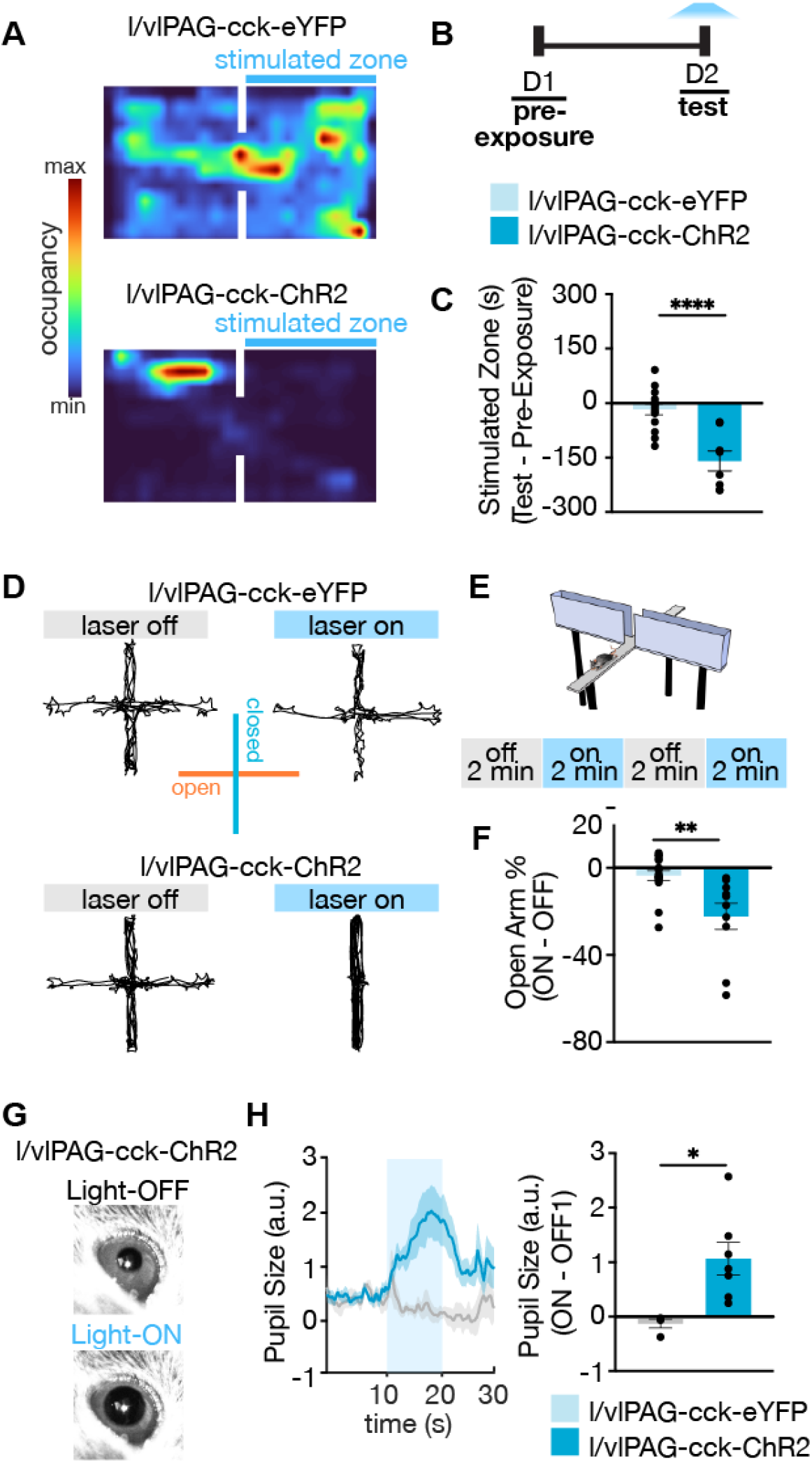
Further characterization of l/vlPAG-cck neurons demonstrates stimulation is aversive, anxiogenic, and induces hallmark sympathetic responses. **(A)** Example spatial map of real-time place preference test (RTPT) depicting min/max occupancy during test of a l/vlPAG-cck-eYFP mouse (top) and l/vlPAG-cck-ChR2 mouse (bottom). Blue light delivery was paired with occupancy of one chamber of the RTPT during Test. **(B)** Timeline for RTPT assay. Each session lasted 10 min. **(C)** Dots represent time spent in the stimulated zone during Test minus time spent in the same zone during Pre-Exposure (without light delivery). Bars are averaged across l/vlPAG-cck-eYFP and l/vlPAG-cck-ChR2-eYFP groups, respectively. Stimulation of l/vlPAG-cck neurons results in avoidance of stimulated zone, compared to control YFP group (eYFP, n = 14; ChR2, n = 8; unpaired t-test, ****p < 0.0001). **(D)** Example locomotion maps in the elevated plus maze (EPM) of l/vlPAG-cck-eYFP (top) and l/vlPAG-cck-ChR2 mice (bottom) during light-off (left) and light-on (right) epochs. **(E)** Stimulation protocol in EPM. **(F)** Dots represent percent of time spent in open arms during light-on epochs normalized by light-off epochs (ON minus OFF) of l/vlPAG-cck-eYFP or l/vlPAG-cck-ChR2 mice. Light delivery to cck-ChR2 mice reduced open-arm occupancy relative to cck-eYFP control mice (eYFP, n = 16; ChR2, n = 10; unpaired t-test, **p < 0.01). **(G)** Example pupil images of a head-fixed l/vlPAG-cck-ChR2 mouse without (top) and with blue-light delivery (bottom). **(H)** Left: Average data showing pupil size during baseline, stimulation, and post-stimulation periods (labeled OFF, ON, and OFF respectively). Each period lasted 10 sec. During stimulation, blue light was delivered to l/vlPAG. Right: Blue light delivery increased pupil size in l/vlPAG-cck-ChR2 but not l/vlPAG-cck-eYFP mice (eYFP, n = 4; ChR2, n = 7; unpaired t-test, *p < 0.05). Mean ± SEM.

### l/vlPAG-cck inhibition delays entry into a dark burrow

Our data show that activation of l/vlPAG-cck neurons is sufficient to induce flight to safety (Figure 3). To determine if these neurons serve a critical role in flight to safety, we next used AAV-mediated, Cre-dependent bilateral expression of the inhibitory opsin archaerhodopsin (Arch) in Cck-ires-cre mice to optically suppress activity of l/vlPAG-cck cells in the Latency to Enter assay (LTE, Figure 5A-B). During Test, green light (562-nm, constant) was delivered to the l/vlPAG in alternating trials and the latency to enter the burrow was measured at the end of each trial (Figure 5B). Though burrow preference was similar across both groups during habituation, light inhibition of l/vlPAG-cck cells increased latency to enter the burrow in Arch mice compared to YFP control mice (Figure 5C-F). In addition to increased l/vlPAG-cck activity playing a sufficient role in inducing flight to safety, these results show l/vlPAG-cck neurons also play a critical role in controlling flight to safety.

**Figure 5.**
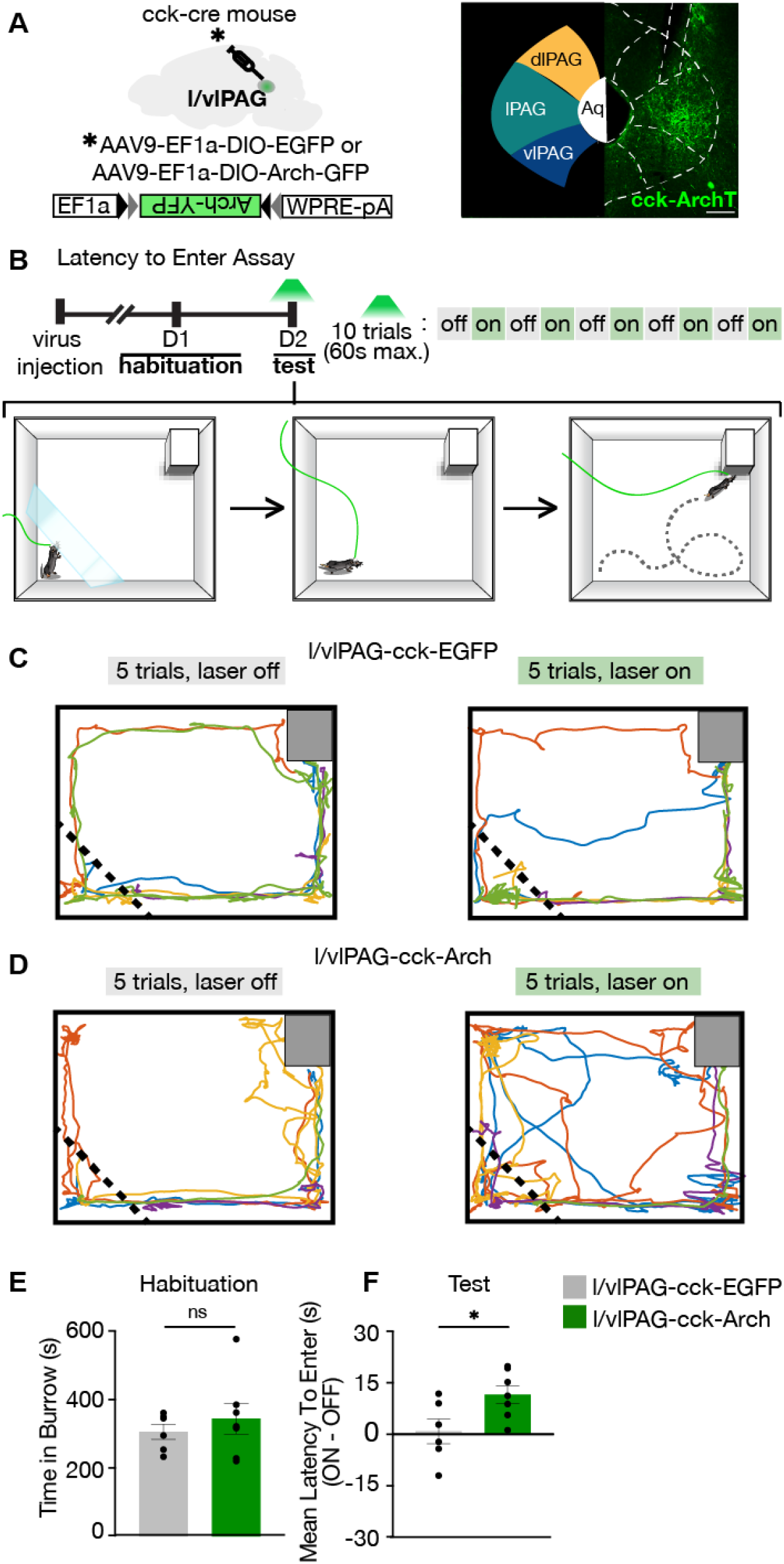
l/vlPAG-cck inhibition delays entry into a dark burrow. **(A)** Left: Strategy for viral expression of cre-dependent GFP or Arch-GFP in l/vlPAG of cck-cre mice. Right: Histology showing Arch-GFP expressed in cck+ cells in the l/vlPAG. Scale bar, 200μm. **(B)** Top: Timeline of Latency to Enter assay. Test consisted of 10 trials, with bilateral green light delivery to l/vlPAG in alternating trials. Bottom: Schematic of assay during test. At the start of trial, mice were confined to a holding zone for 15s with a transparent barrier. When the barrier was removed, mice were free to explore the arena. Trial ended upon burrow entry or 60s had passed. **(C)** Example locomotion maps of five trials without (left) and with (right) green light delivery in a cck-GFP mouse. **(D)** Same as (C) but in a cck-Arch mouse. **(E)** No difference in burrow occupancy during 10-min habituation between cck-GFP and cck-Arch mice (GFP, n = 6; Arch, n = 7; unpaired t-test). **(F)** Green light delivery to l/vlPAG increased latency to enter burrow in cck-Arch mice compared to cck-GFP mice. Each dot represents average latency during 5 light-on trials minus average latency of 5 light-off trials (GFP, n = 6; Arch, n = 7; unpaired t-test, *p < 0.05). Mean ± SEM.

### l/vlPAG-cck stimulation increases avoidance of a live predator

Our data show that l/vlPAG activity is sufficient and necessary for flight to safety in the LTE, a low-threat environment in which perceived danger is diffuse and uncertain (La-Vu et al., 2020). However, it is still unknown if the l/vlPAG-cck population is involved in flight to safety in the presence of a well-defined, proximal threat such as a live predator. To address this question, we optogenetically activated l/vlPAG-cck cells while introducing mice to the Live Predator assay (Figure 6A). In this assay, mice are placed in an elongated rectangular arena that contains an awake rat restrained to one end by a harness (Reis et al., 2021; Wang et al., 2021). Rats are natural predators of mice and mice exhibit robust defensive reactions during exposure to a live rat but not a similarly-shaped toy rat (Figure 6B, (Wang et al., 2021). As the chamber does not contain a barrier and mice can freely roam the entire arena, the Live Predator assay elicits a naturalistic and diverse repertoire of defensive responses (Reis et al., 2021; Wang et al., 2021). We hypothesized that if l/vlPAG-cck cells serve a role in flight to safety from predatory threat, then activation of these cells would exacerbate active avoidance of a live rat. To test this hypothesis, we delivered blue light (473-nm, 5-ms, 20-Hz) to the l/vlPAG of mice expressing ChR2-YFP or YFP in l/vlPAG-cck cells in the Live Predator assay (Figure 6C). Light was delivered in alternating 2-min epochs (Figure 6D-E). Light activation of l/vlPAG-cck cells reduced time spent in the threat zone and increased distance from the live rat (Figure 6F-G). Mice exhibit increased stretch-attend postures during exposure to predatory rats (Wang et al., 2021). This measure was reduced as a result of optogenetic activation, demonstrating that not all defensive behaviors are promoted by cck cell activation (Figure 6I). Optogenetic activation of cck cells also induced a trend toward reduced number of approaches toward the rat, and did not alter freezing or locomotion (Figure 6H, J-K). Escape velocity can be an informative measure of threat avoidance; however, ChR2 mice did not exhibit enough escapes during light activation to compute this measure as they did not consistently approach the rat (see representative exploration track in Figure 6E, lower row), which decreased escapes from the rat as escapes cannot occur without prior approach. These results demonstrate activation of l/vlPAG-cck cells selectively enhanced avoidance of a live predator without altering freezing. Importantly, these same results were not observed during cck activation during exposure to a control toy rat (Figure 6, figure supplement 1). Activating cck cells in this condition induced the same type of thigmotaxis seen during cck activation in the open field (Figur 2G). In the presence of the toy rat, thigmotaxis was uniformly induced throughout the environment periphery, both near and far away from the toy rat (see representative exploration track in Figure 6, figure supplement 1A). Thus, in the presence of the toy rat, cck activation induced avoidance of open spaces, rather than avoidance of the toy rat. In contrast, in the presence of the rat, activation of cck cells induced thigmotaxis only in the corners furthest away from the live rat (Figure 6E). These data show that l/vlPAG cck cell activation increases avoidance of a live predator, but not of a control safe toy rat, showing that these cells may serve to minimize threat exposure by directing exploration towards safer regions within an environment.

**Figure 6.**
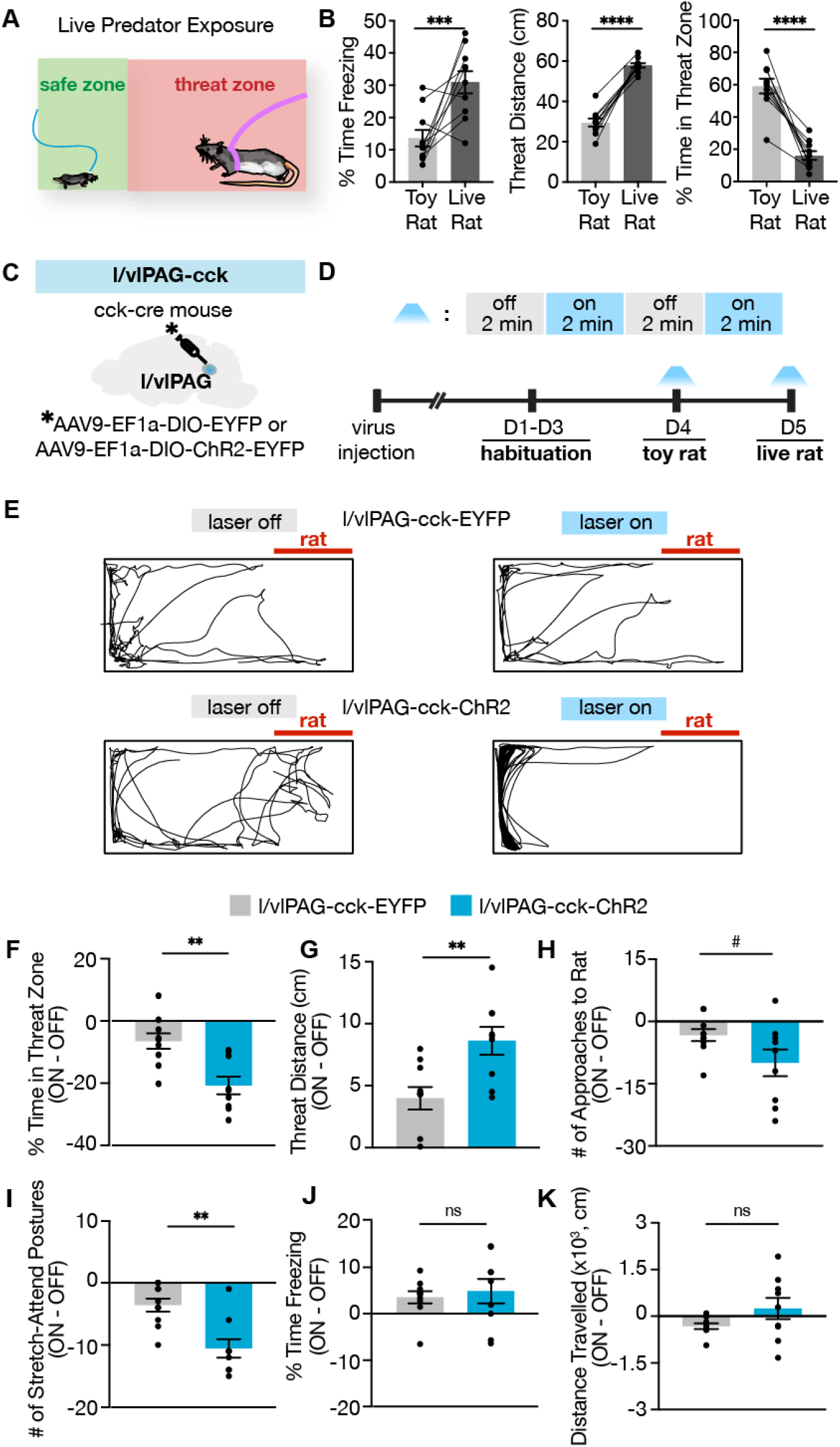
l/vlPAG-cck activation robustly enhances active avoidance from a live predator without altering freezing. **(A)** Schematic of Live Predator Exposure assay. Mice are placed in a long rectangular chamber (70 × 25 × 30 cm) containing an awake rat restrained with a harness to one end. The chamber does not contain a barrier and mice can move freely. The area containing the rat is considered a ‘threat zone’ and the area furthest from the rat is considered a ‘safe zone.’ **(B)** Exposure to a live rat increased freezing and threat distance while reducing time in threat zone compared to exposure to a toy rat (n = 10, paired t-test, ***p < 0.001, ****p < 0.0001). **(C)** EYFP or ChR2-eYFP was expressed in l/vlPAG-cck cells and a fiber-optic cannula was implanted over the l/vlPAG. **(D)** Timeline of Live Predator assay. Blue light was delivered in alternating 2-min epochs during Toy Rat exposure and Live Rat exposure. **(E)** Example locomotion maps during laser-off (left) and laser-on (right) epochs of an eYFP mouse (top) and ChR2-eYFP mouse (bottom). **(F-K)** Optogenetic stimulation of l/vlPAG-cck cells reduced time in threat zone (**F**), increased threat distance (**G**), and reduced stretch-attend postures **(I**). Number of approaches to the rat trended toward significance (H, #p = 0.066). Freezing (**J**) and distance traveled (**K**) were unaffected. (eYFP, n = 10; ChR2, n = 9; unpaired t-test, ns = not significant, **p < 0.01). Mean ± SEM.

### l/vlPAG-cck inhibition decreases avoidance of a live predator

To evaluate the necessity of l/vlPAG-cck cells for flight to the safer region in a high-threat environment, we bilaterally expressed Cre-dependent inhibitory hM4di-mCherry in the l/vlPAG of *Cck-ires-cre* mice to chemogenetically inhibit l/vlPAG-cck cells in the Live Predator assay (Figure 7A). Mice expressing mCherry alone served as controls. A chemogenetic approach in this setting was beneficial because it enabled neuronal inhibition across a 10-min exposure without constant laser delivery, as prolonged laser stimulation may induce tissue heating, among other issues ((Stujenske et al., 2015)). Both hM4di and mCherry-only mice were injected with clozapine-N-oxide (CNO) or saline prior to two exposures to a toy rat and two exposures to an awake, live rat on separate, sequential days (Figure 7B-C). Injections occurred 40 min prior to exposure and the order of drug delivery was counterbalanced across groups. All metrics were calculated as behavior exhibited following saline administration subtracted from behavior exhibited following CNO administration (CNO - SAL). We found that l/vlPAG-cck inhibition significantly increased time spent in the threat zone, increased the number of approaches toward the rat, and reduced escape velocity from the rat (Figure 7D, F, H). L/vlPAG-cck inhibition also induced a trend towards decreased distance from the rat (Figure 7E). Inhibition did not alter approach velocity, stretch-attend postures, freezing, or distance traveled (Figure 7G, I-K). Importantly, l/vlPAG-cck inhibition did not affect avoidance from a toy rat (Figure 7, figure supplement 1), indicating effects of inhibition are specific to a live predator. Inhibition of cck cells did not affect pain response latency during exposure to a heated plate assay or acquisition of conditioned fear (Figure 7, figure supplement 2), demonstrating that these cells do not affect learned defensive responses and that they do not control other PAG functions such as analgesia (Samineni et al., 2017). Together, these results demonstrate a critical role for the l/vlPAG-cck population in selectively promoting flight, and not other defensive responses.

**Figure 7.**
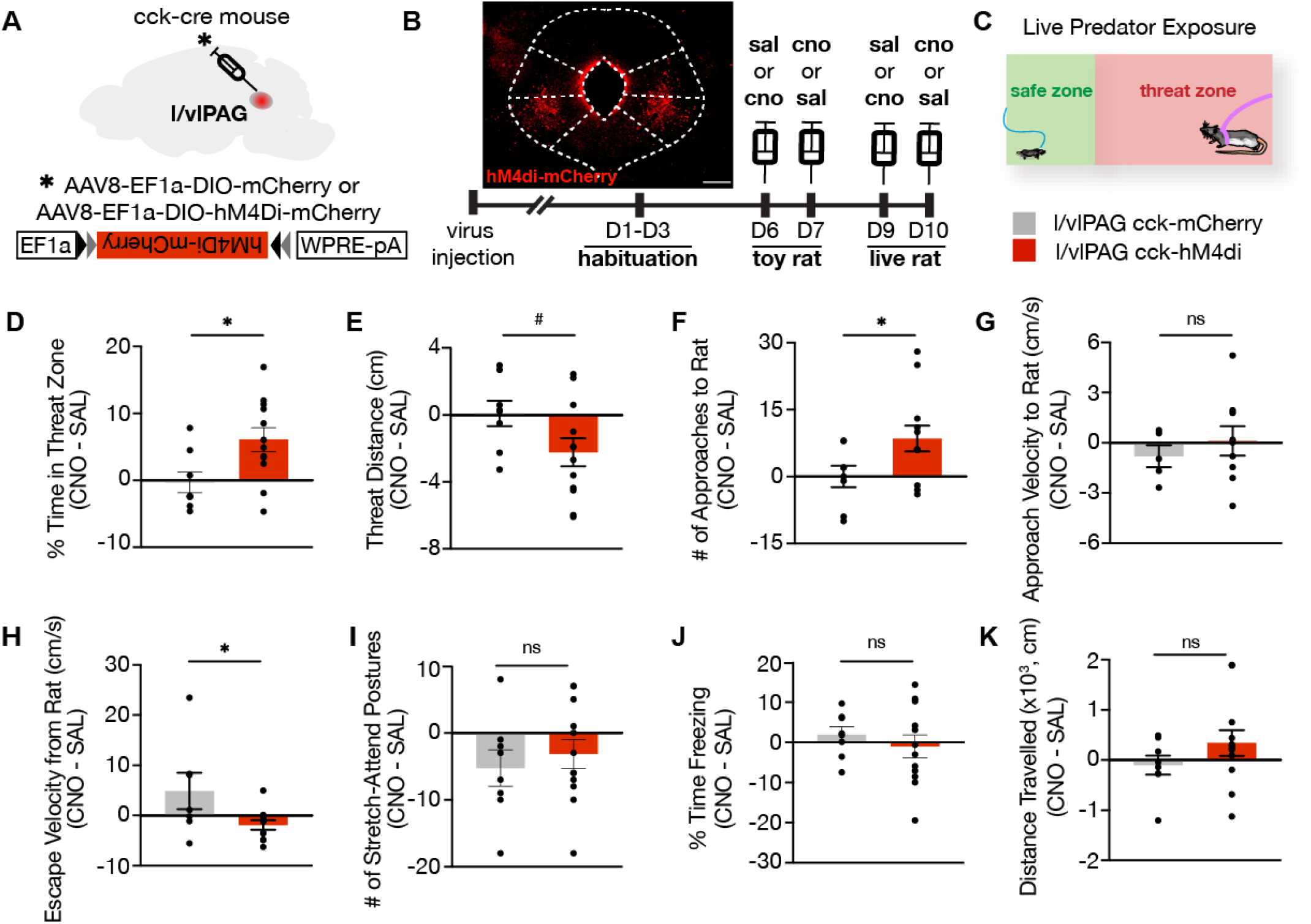
l/vlPAG-cck inhibition increases time spent near a live predator and reduces escape vigor without altering freezing. **(A)** Viral strategy for bilateral expression of inhibitory designer receptor hM4Di-mCherry or control mCherry-only in cck cells in the l/vlPAG. **(B)** Top-left: Expression of hM4Di-mCherry in l/vlPAG-cck cells. Scale bar, 200μm. Bottom: Timeline for DREADD experiments. Saline or clozapine-N-oxide (CNO, 10mg/kg) was administered 40 min prior to exposure. **(C)** Live Predator exposure schematic. Mice were placed in the presence of an awake rat restrained to one end of a chamber. Each exposure lasted 10 min. **(D-K)** Chemogenetic inhibition of l/vlPAG cck cells increased time spent in threat zone (**D**), increased number of approaches toward the rat (**F**), and reduced escape velocity from the rat (**H**). Threat distance trended toward significance with CCK inhibition (E, #p = 0.069). Approach velocity (**G**), stretch-attend postures (**I**), freezing (**J**), and distance traveled (**K**) were unaltered with inhibition (**D-F, I-K**: mCherry, n = 8; hM4Di, n = 12; G: mCherry, n = 5; hM4Di, n = 9; **H**: mCherry, n = 7; hM4Di, n = 11; unpaired t-test, *p < 0.05). Mean ± SEM.

### l/vlPAG-syn cells are more active near threat, while l/vlPAG-cck cells are more active far from threat

Numerous prior reports have consistently shown that PAG cells are activated by proximity to danger (Deng et al., 2016; Evans et al., 2018; Mobbs et al., 2010, 2007; Reis et al., 2021; Watson et al., 2016). We next sought to observe endogenous l/vlPAG-cck activity under low-threat and high-threat conditions. We performed *in vivo* fiber photometry recordings of synapsin and cck-expressing neurons in the l/vlPAG in the elevated plus maze (EPM) and live predator exposure assays (Figure 8A-C).

**Figure 8.**
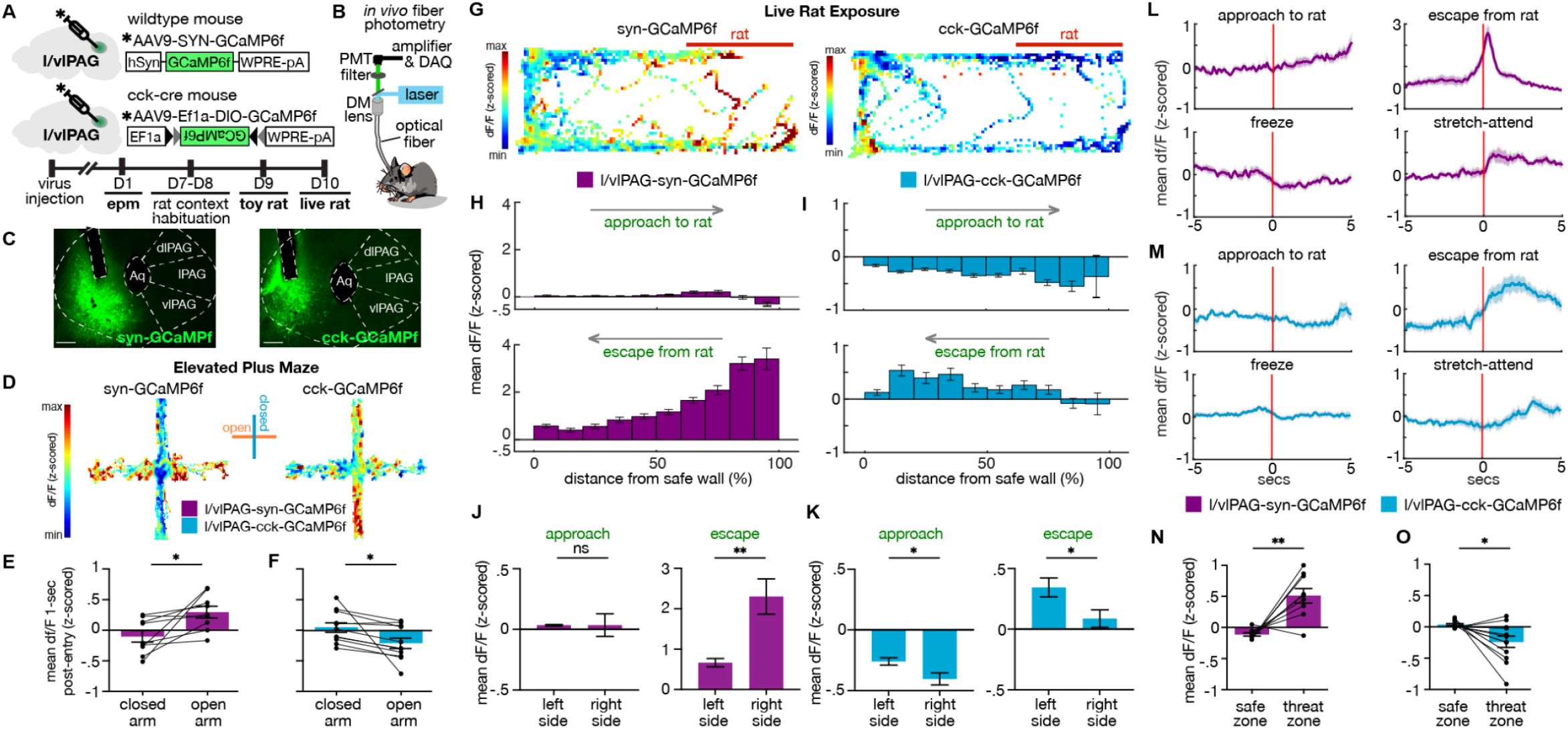
l/vlPAG-syn cells are more active near threat, while l/vlPAG-cck cells are more active far from threat. **(A)** Top, viral strategy for synapsin-specific and cck-specific GCaMP6f expression in l/vlPAG. Bottom, timeline for *in vivo* photometry recordings. **(B)** Fiber photometry recording set-up. **(C)** Histology of GCaMP6f expression in synapsin-specific (left) and cck-specific (right) cells in the l/vlPAG. Scale bar, 200μm. **(D)** Example heat maps showing z-scored dF/F in mice expressing synapsin-specific GCaMP6f (left) or cck-specific GCaMP6f (right) in l/vlPAG in an elevated plus maze. Vertical arms of heat maps represent closed arms. **(E-F)** Mean dF/F (z-scored) one-second after arm entry in syn-GCaMP6f (**E**) and cck-GCaMP6f mice (**F**). Mean dF/F 1-sec post-entry into the open arms is greater than into closed arms in syn-GCaMP6f mice (**E**), whereas mean dF/F 1-sec post-entry into the open arms is lower than into the closed arms for cck-GCaMP6 mice (syn, n = 9; cck, n = 11; paired t-test, *p < 0.05). **(G)** Example heat maps showing z-scored dF/F in syn-GCaMP6f (left) or cck-GCaMP6f (right) during Live Rat exposure. Rat was confined to the right of the map, as indicated by the red bar. **(H-I)** Mean dF/F (z-scored) during approaches toward the rat (top) or escapes from the rat (bottom) within ten spatial bins of varying distance from the safe wall of syn-GCaMP6f (**H**) or cck-GCaMP6f (**I**) mice (syn, n = 9; cck, n = 13; syn-approach, n = 6744 samples; syn-escape, n = 2150 samples; cck-approach, n = 7170 samples; cck-escape, n = 2088 samples). **(J-K)** Comparisons of mean dF/F in five left (close to safe wall) bins to five right (further from safe wall) bins in H-I, respectively. (**J**) Syn-GCaMP6f activity was not modulated during approaches to the rat but was robustly modulated during escape from the rat, reducing as mice gain distance from rat. (**K**) Cck-GCaMP6f activity was modulated during approach, decreasing as mice moved closer to the rat. Cck activity was also modulated during escape, increasing as mice gained distance from rat (left side, n = 5 bins; right side, n = 5 bins; unpaired t-test, *p < 0.05, **p < 0.01). **(L-M)** Mean dF/F (z-scored) 5-sec before and after approaches, escapes, freeze bouts, and stretch-attend postures in syn-GCaMP6f (**L**) and cck-GCaMP6f (**M**) populations (syn, n = 9; cck, n = 13 for freeze, n = 12 for other behaviors). **(N-O)** Mean dF/F (z-scored) in the safe zone (one-third of assay near safe wall) and threat zone (two-thirds of assay distal from safe wall) in syn-GCaMP6 (**N**) and cck-GCaMP6 (**O**) mice. Pan-neuronal l/vlPAG activity was increased in the threat zone compared to the safe zone, whereas cck-specific activity was decreased in the threat zone compared to the safe zone (syn, n = 9; cck, n = 12; paired t-test, *p < 0.05, **p < 0.01). Mean ± SEM.

These assays offer a safety gradient that allow us to assess how population activity is spatially modulated by threat proximity. Recording of syn-expressing l/vlPAG cells will inform whether activity patterns in cck-only population recordings are cell type-specific or region-specific. We found that l/vlPAG-syn and l/vlPAG-cck population activity was differentially modulated by EPM arms (Figure 8D). Specifically, mean df/F of l/vlPAG-syn cells increased after open-arm entry compared to closed arm entry (Figure 8E). In contrast, cck activity was greater following closed arm entry compared to open arm entry (Figure 8F). In a low-threat setting such as the EPM, l/vlPAG-cck activity diverged from syn-expressing l/vlPAG cell activity.

To determine if this feature might extend to a high-threat situation, we next performed photometry recordings of l/vlPAG pan-neuronal and cck-only populations during live rat exposure (Figure 8G). Mean df/F within spatial bins of varying distance from the safe wall shows l/vlPAG-syn activity was not differentially modulated during approaches toward rat but was significantly altered during escapes, exhibiting increased activity at initiation of escape and ramping down as mice gained distance from the rat (Figure 8H,J). This pattern is consistent with previous reports coupling increased PAG activity with threat proximity and escape initiation (Deng et al., 2016; Evans et al., 2018; Reis et al., 2021; Watson et al., 2016). Contrarily, l/vlPAG-cck activity was modulated during both approach and escape, ramping down as mice approached the rat and ramping up as mice escaped from the rat (Figure 8 I,K). l/vlPAG-syn activity was tightly time-locked with escapes, ramping up prior to and peaking soon after escape initiation; l/vlPAG-cck activity also ramped up prior to escape but exhibited prolonged increased activation after initiation (Figure 8L-M). Our optogenetic experiments showed that increasing activity in l/vlPAG-syn cells robustly induced freezing; however we did not observe increased l/vlPAG-syn population activity during freeze bouts during predator exposure (Figure 8L). Finally, overall l/vlPAG-syn activity was greater in the threat zone than safe zone while l/vlPAG-cck activity was decreased in the threat zone relative to the safe zone (Figure 8N-O).

Importantly, these effects were not due to a correlation between speed and df/F (Figure 8, figure supplement 1) and were not observed during exposure to a toy rat (Figure 8, figure supplement 2). Together, these results suggest that though the syn-expressing l/vlPAG population was particularly attuned to more threatening aspects of both the EPM and live predator exposure, cck population activity was negatively modulated by threat proximity and appears involved in the transition from danger to safety.

## Discussion

Our study identifies a small, sparse, genetically-defined subset of cells within the PAG, cholecystokinin-expressing neurons, that robustly controls flight to safety in multiple contexts. We show this feature cannot be induced with pan-neuronal activation of the same location. L/vlPAG-cck cells can bidirectionally control flight to safety in low- and high-threat conditions without altering freezing. In contrast, pan-neuronal activation of lPAG and vlPAG drives freezing (Figure 2J), as consistently shown previously (Assareh et al., 2016; Bittencourt et al., 2005; Tovote et al., 2016; Yu et al., 2021). Endogenous Ca^2+^ recordings show that while pan-neuronal l/vlPAG activity is increased with threat proximity consistent with prior reports in the PAG (Deng et al., 2016; Evans et al., 2018; Mobbs et al., 2010, 2007; Reis et al., 2021; Watson et al., 2016), l/vlPAG-cck activity is decreased with threat proximity and is attuned to transition from danger to safety.

Importantly, though our strategy of synapsin-specific transfection does not exclude cck+ cells, we show cck+ cells are a small, sparse population, making up only about 5% of l/vlPAG neurons (Figure 1), and are unlikely to be significantly driving fluorescence in pan-neuronal photometry recordings.

One inconsistency in our data findings is a lack of increase in l/vlPAG-syn fluorescence during freeze bouts in the predator assay, despite ChR2 activation of l/vlPAG-syn cells driving robust freezing. This may be due to several factors including ChR2 activation may stimulate more ventral cells than are being recorded using fiber photometry. Another possibility is that ChR2 activation may drive activity in some few cells that are responsible for driving the robust freezing observed, and the fluorescence of these sparse cells were washed out in Ca^2+^ recordings. Finally, it is possible that l/vlPAG activity reflected complex population dynamics related to the heightened behavioral state induced by high-threat predator exposure, resulting in endogenous activity that is more complex than is elicited with artificial activation in a low-threat environment.

### Role of cck cells in the l/vlPAG

Long-standing evidence links increased PAG activity with greater threat exposure. Increased Fos expression has been reported in the rodent PAG following predator exposure (Aguiar and Guimarães, 2009; Canteras and Goto, 1999; Mendes-Gomes et al., 2020). In addition, pharmacological blockade of NMDA receptors in the dorsal or ventrolateral PAG increases open arm exploration in the EPM (Guimarães et al., 1991); (Molchanov and Guimarães, 2002). Single unit recordings show dPAG and vPAG units display significant increases in firing rate after exposure to cat odor (Watson et al., 2016). Moreover, the dPAG is sensitive to sensory aspects of threat distance and intensity, displaying increased activity with greater threat proximity (Deng et al., 2016). In humans, PAG activity is also positively correlated with threat imminence (Mobbs et al., 2010, 2007). Within the dPAG, glutamatergic neurons are key for escape initiation and vigor, and dPAG flight-related cells exhibited prominent firing early during flight and declined as mice fled further from a predator (Deng et al., 2016; Evans et al., 2018).

Similarly, our *in vivo* recordings of pan-neuronal l/vlPAG cells showed enhanced activity after EPM open arm entry compared to closed arm entry. During predator exposure, l/vlPAG activity was also closely time-locked to initiation of escape from threat and ramped down as mice gained distance from threat. Together, our findings suggest a role of l/vlPAG neurons in the motor aspect of escape initiation and representation of threat during flight, concordant with previous reports of the PAG (Assareh et al., 2016; Bittencourt et al., 2005; Tovote et al., 2016; Yu et al., 2021).

In contrast, a sparse population of glutamatergic cck+ cells in the same region were more active after entry into closed arms than open arms of the EPM. Like synapsin-expressing cells during predator exposure, cck+ cells also ramped up upon initiation of escape; however, they showed prolonged enhanced activity post-escape and further ramped up as mice gained distance from threat. Thus, unlike pan-neuronal cells that appear attuned to the motor action of escape initiation, our findings suggest cck+ cells are more involved in the transition from high threat to relative safety during escapes.

The lateral and ventrolateral PAG have been shown to control freezing. Prior optogenetic stimulation of lateral PAG caused freezing at low stimulation intensity and escape in high intensity (Assareh et al., 2016). Similarly to our results of pan-neuronal excitation of l/vlPAG cells, optogenetic excitation of lateral PAG vGlut2+ cells produces robust freezing (Yu et al., 2021). Electrical lateral PAG excitation also caused freezing (Bittencourt et al., 2005). Glutamatergic neurons in the ventrolateral PAG, at the control of a local disinhibitory circuit, powerfully mediate freezing (Tovote et al., 2016). Furthermore, lesions to the ventral PAG curtails freezing during predator exposure and conditioned fear (de Andrade Rufino et al., 2019; Fanselow et al., 1995). However, we show that activation of cck+ l/vlPAG cells reliably causes flight to relative safety in multiple assays (open field, LTE, EPM, and predator). Interestingly, the flight pattern observed from cck+ l/vlPAG activation is also characteristically different from escape canonically described in PAG studies, as they are void of robust vertical jumping as seen in dlPAG activation in this study (Figure 2I) and prior work (Ullah et al., 2015). Our findings suggest the region’s involvement in defensive behaviors may be broader than just freezing, containing a cck+ population that powerfully drives flight to safety and is preferentially active during transition from danger to safety.

Importantly, our work draws attention to two distinct types of escape: flight to escape the environment, as seen by jumps induced by broad pan-neuronal activation of dlPAG cells (Figure 2I), and flight to safer regions within the environment, as shown by activation of l/vlPAG cck cells (Figure 6). Our findings show that cck cells are specifically involved in flight to safer regions within an environment, as protean escape jumps were not observed in any mouse. Further studies are needed to identify how l/vlPAG cck cells affect downstream targets to produce escape, and also to investigate which inputs to these cells are necessary for activating them.

### Complementing columnar functional organization

Columnar organization in the PAG is supported by functional and anatomical similarities along the rostrocaudal axis. Broad activation of the dorsolateral column induces flight, hypertension, and tachycardia, while activation of the ventrolateral column induces freezing, hypotension and bradycardia (Keay and Bandler, 2015). Neurotransmitter and receptor expression profiles and afferent/efferent connections are also generally (but not always) homogeneous within a single column along the anteroposterior axis (Silva and McNaughton, 2019).

However, to achieve a more complete understanding of PAG function, there may be other considerations in addition to columnar organization. There are exceptions to homogeneity along the anterior-posterior axis of columnar boundaries in PAG afferent and efferent connectivity. For example, adrenergic and noradrenergic medullary afferents preferentially target the rostral vlPAG and the central amygdala receives input from cells highly concentrated in the caudal but not rostral vlPAG (Silva and McNaughton, 2019). Furthermore, there are genetic markers that do not span an entire column; GABA-immunopositive cells are more prevalent in caudal than rostral cat vlPAG (Barbaresi, 2005). Tachykinin-1 (i.e., tac1), which is a marker of substance P-expressing cells, broadly spans multiple columns rostrally but concentrates in dorsolateral and ventrolateral columns caudally in the rat (Liu and Swenberg, 1988). Expression of rat endocannabinoid and glycine receptors also vary rostrocaudally, becoming more present in caudal PAG (Araki et al., 1988; Herkenham et al., 1991; Silva and McNaughton, 2019). It is likely that exploration of these genetically-defined PAG populations will reveal novel insights. Indeed, inhibition of lPAG Vgat and lPAG Vglut2 neurons impair the chase and attack of prey, respectively (Yu et al., 2021), and glutamatergic vlPAG neurons project to the medulla to control freezing (Tovote et al., 2016). Moreover, l/vlPAG tac1 cells have been shown to specifically control itching behavior (Gao et al., 2019), further supporting the value of investigating sparse genetically-defined PAG populations.

Recent work has shown that examination of genetic diversity can unveil deep and novel understanding even in well-studied regions such as the amygdala. Accordingly, work from numerous groups dissecting glutamatergic basolateral amygdala cells based on genetic markers and projection targets has revealed the region’s complex control of anxiety and valence processing (Felix-Ortiz et al., 2013; Kim et al., 2016; Tye et al., 2011).

In this study, using a genetic approach to target cck+ cells, we uncovered a role for these cells in flight to safe regions within the present environment. Cck-expressing cells are one among many largely uncharacterized genetically-defined populations in the PAG (Yin et al., 2014), and here we outline a framework to assess how a single cell type may contribute to the vast constellation of behaviors controlled by the PAG. These results highlight that the molecular identity of PAG cells can lend key insight into functional motifs that govern how the PAG produces defensive responses and may serve as an additional axis of functional organization complementing the well-established anatomical columnar PAG divisions.

## Acknowledgements

We were supported by the NIMH (R00 MH106649 and R01 MH119089 to A.A. and F31 MH121050-01A1 to M.L-V.); the Achievement Rewards for College Scientists Foundation, Los Angeles Chapter (to M.L-V.); the Brain and Behavior Research Foundation (22663, 27654, and 27780 to A.A., F.M.C.V.R., and W.W. respectively); the NSF (NSF-GRFP DGE-1650604 to P.S.); the UCLA Affiliates fellowship (to P.S.); the Hellman Foundation (to A.A.); F.M.C.V.R. was supported by FAPESP grants 2015/23092-3 and 2017/08668-1. We thank Profs. S. Correa, K. M. Wassum, and M. Fanselow for helpful discussions. We thank H.T. Blair and K.M. Wassum for providing rats.

## Figures

**Figure 6, figure supplement 1.**
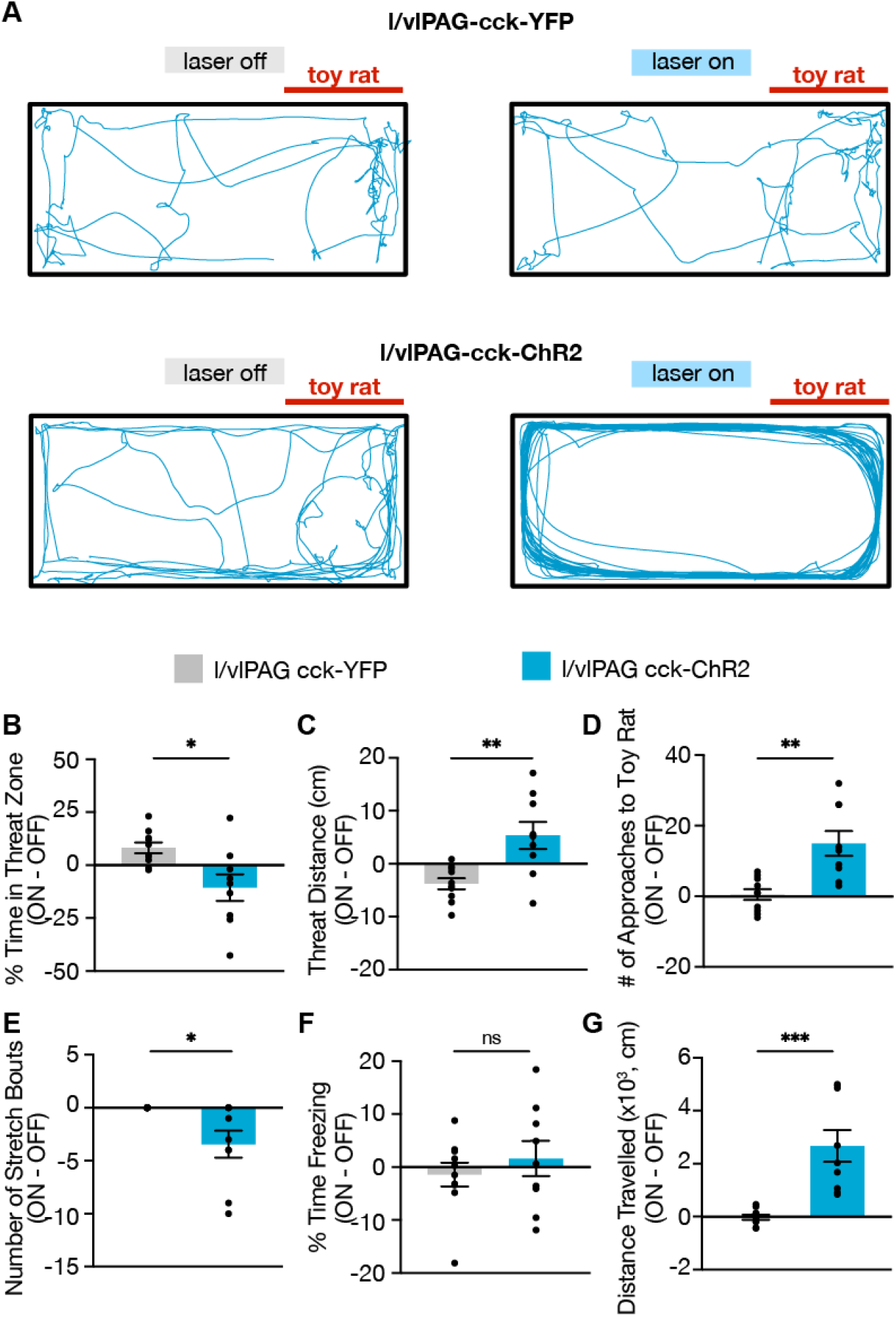
Optogenetic activation of l/vlPAG-cck neurons during control toy rat exposure. **(A)** Example locomotion maps during laser-off (left) and laser-on (right) epochs of an l/vlPAG-cck-eYFP mouse (top) and an l/vlPAG-cck-ChR2-eYFP mouse (bottom). Stimulation induced robust traversal of all four corners of the enclosure (bottom-right). **(B-G)** Optogenetic stimulation of l/vlPAG-cck cells reduced time spent in threat zone (**B**) increased threat distance (**C**), increased number of approaches to the toy rat (**D**), reduced stretch bouts (**E**), and increased distance traveled (**G**). Stimulation did not alter freezing (**F**) (eYFP, n = 10; ChR2, n = 9; unpaired t-test, ns = not significant, *p < 0.05, **p < 0.01, ***p < 0.001). Mean ± SEM.

**Figure 7, figure supplement 1.**
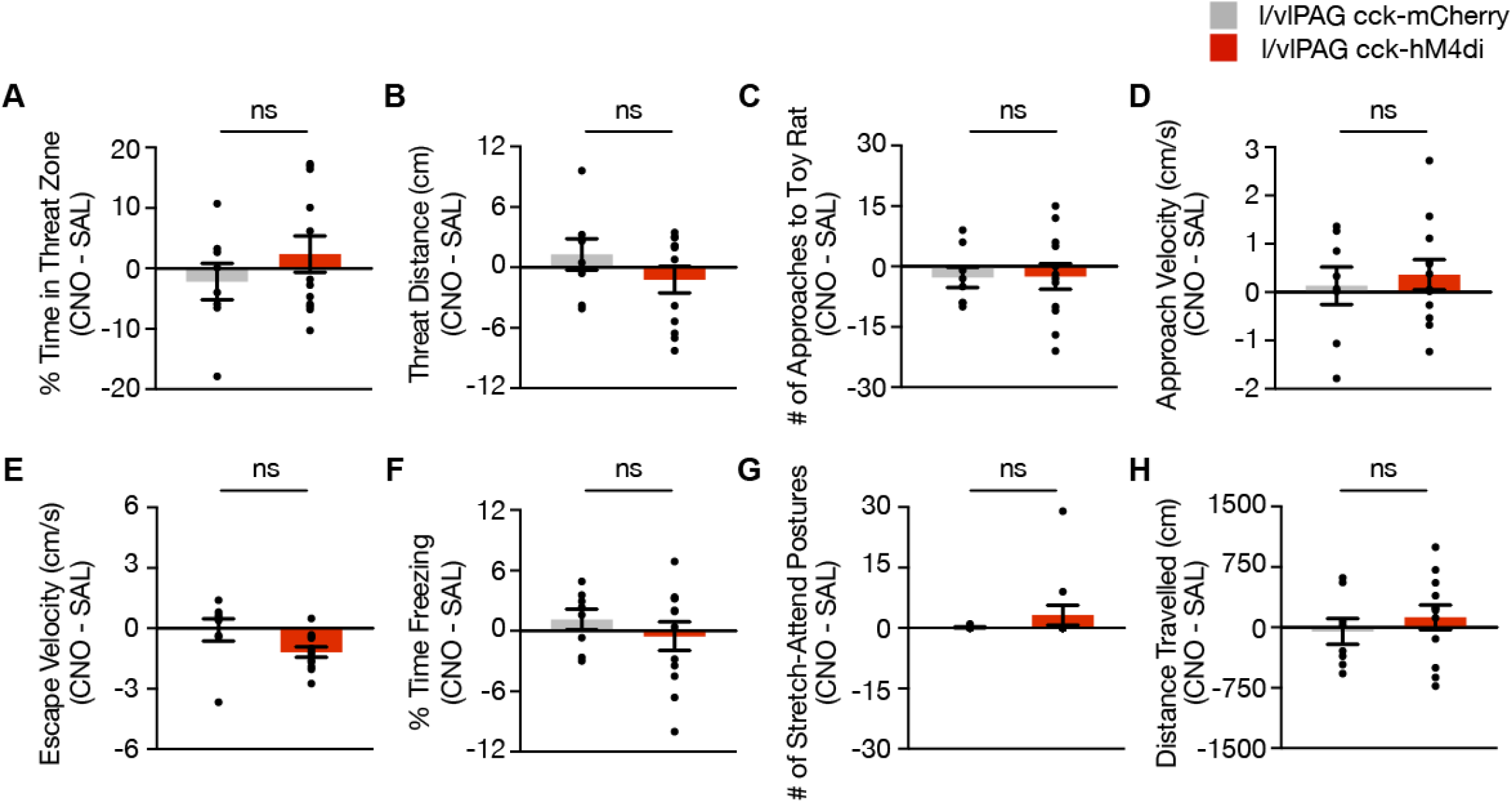
Chemogenetic inhibition of l/vlPAG-cck population using DREADDs during exposure to a control toy rat. **(A-H)** Chemogenetic inhibition of l/vlPAG-cck cells during exposure to toy rat (mCherry, n = 8; hM4Di, n = 12; unpaired t-test). Mean ± SEM.

**Figure 7, figure supplement 2.**
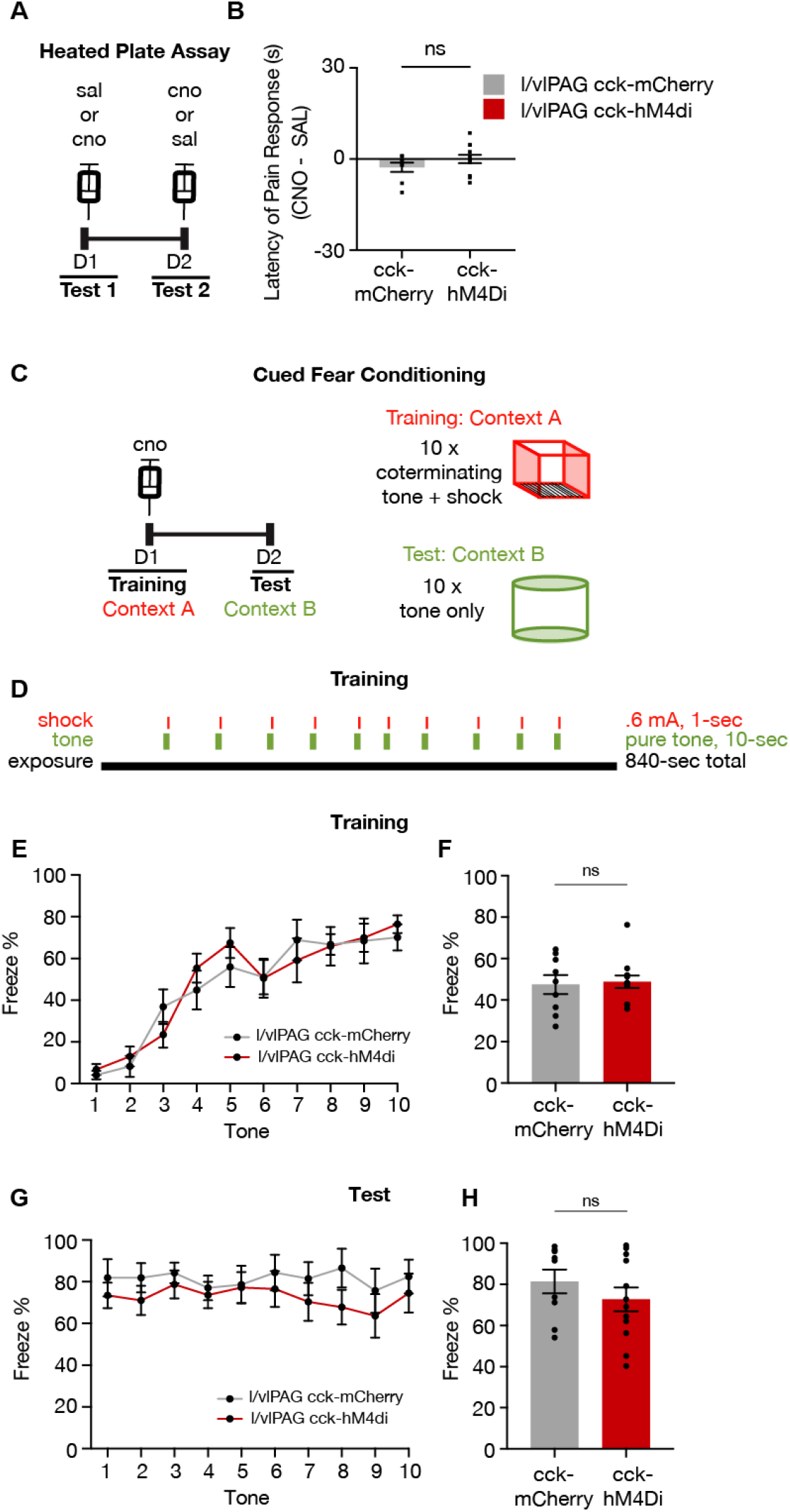
Inhibition of l/vlPAG-cck neurons does not alter pain response latency or acquisition of learned fear. **(A)** Timeline of heated plate assay with chemogenetic inhibition of l/vlPAG-cck cells. Order of drugs was counterbalanced across groups. **(B)** Inhibition of l/vlPAG-cck cells does not alter latency of pain response (CNO minus saline; mCherry, n = 8; hM4Di, n = 12; unpaired t-test). Mean ± SEM. **(C)** Timeline of cued fear conditioning with l/vlPAG-cck chemogenetic inhibition. All mice received CNO prior to Training. **(D)** Schedule of tone and shock presentations during Training. Trial was 14-min with ten co-terminating 10-sec tones and 1-sec shocks. **(E)** Mean freezing of cck-mCherry and cck-hM4Di mice during 10-sec tones during Training (mCherry, n = 9; hM4Di, n = 12). **(F)** No difference in freezing during Training between groups (mCherry, n = 9; hM4Di, n = 12; unpaired t-test). **(G)** Same as E but during Test (mCherry, n = 9; hM4Di, n = 12). **(H)** Same as F but during Test (mCherry, n = 9; hM4Di, n = 12; unpaired t-test).

**Figure 8, figure supplement 1.**
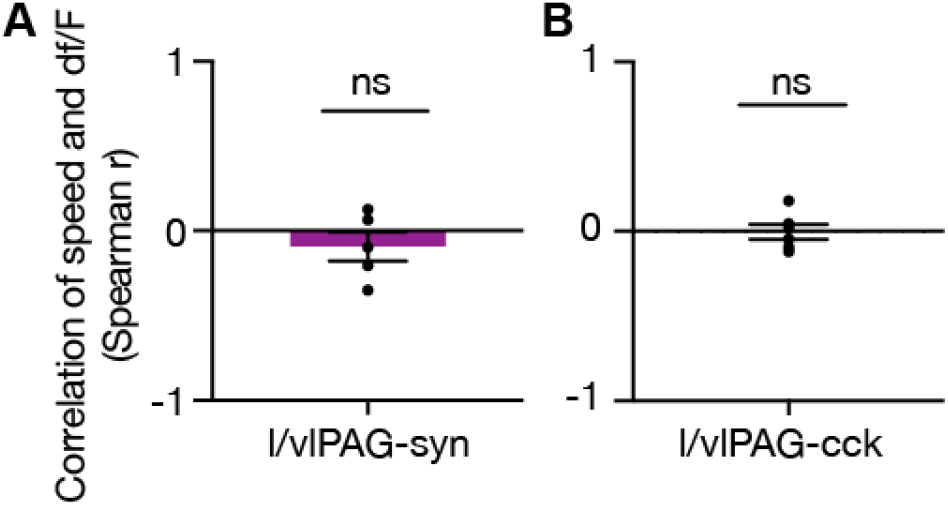
No correlation between speed and df/F. **(A-B)** Spearman correlation of speed and fiber photometry df/F of (**A**) l/vlPAG-synapsin or (**B**) l/vlPAG-cck population during exposure to toy rat (syn, n = 5; cck, n = 6; one-sample t-test). Mean ± SEM.

**Figure 8, figure supplement 2.**
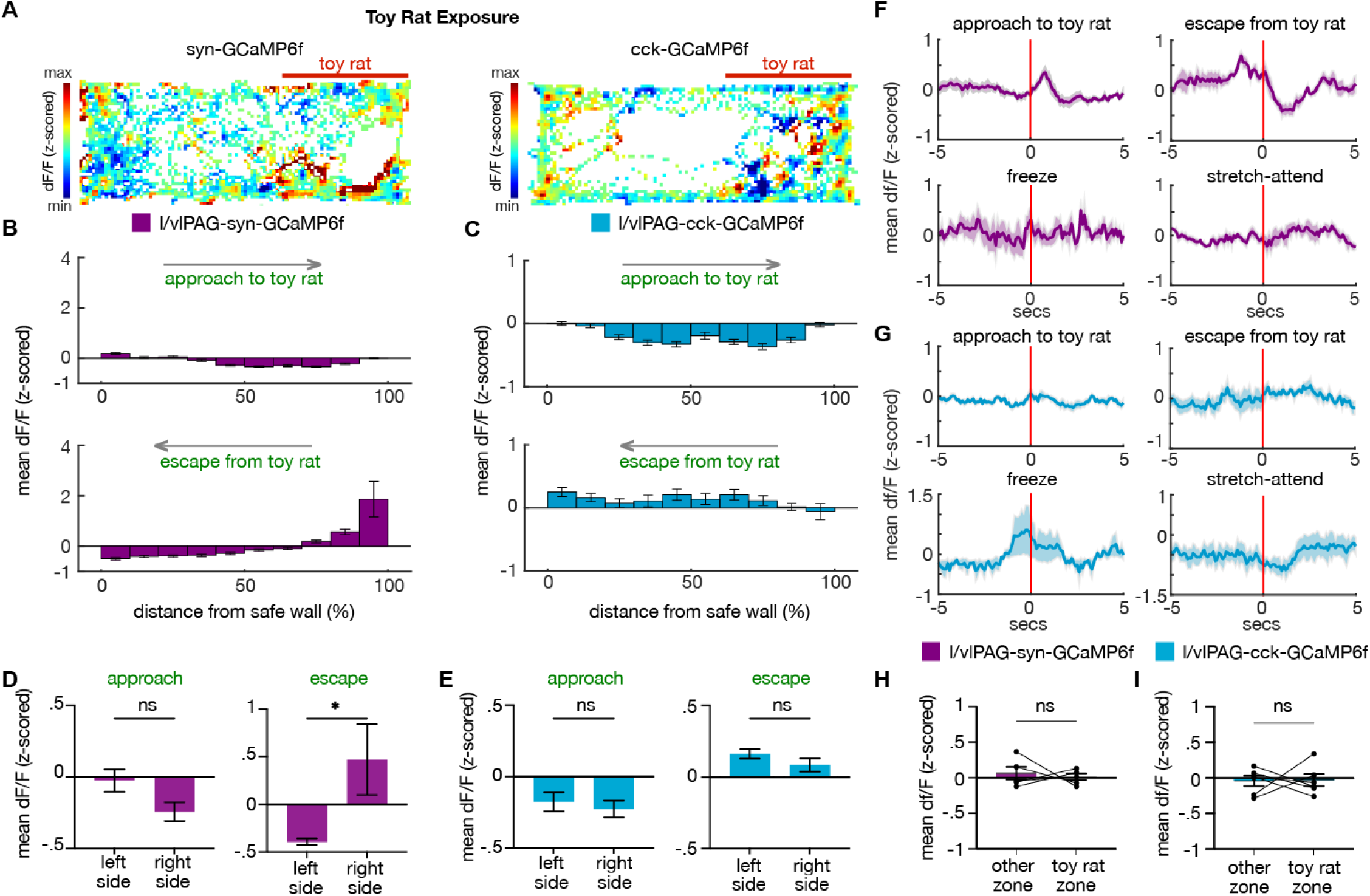
l/vlPAG-syn- and cck-expressing cells do not encode proximity to a control toy rat. **(A)** Example heat maps showing z-scored dF/F in syn-GCaMP6f (left) or cck-GCaMP6f (right) during exposure to a control toy rat. The toy rat was confined to the right of the map, as indicated by the red bar. **(B-C)** Mean dF/F (z-scored) during approaches toward the toy rat (top) or escapes from the toy rat (bottom) within ten spatial bins of varying distance from the safe wall of syn-GCaMP6f (B) and cck-GCaMP6f (C) mice (syn, n = 5; cck, n = 6; syn-approach, n = 7361 samples; syn-escape, n = 1884 samples; cck-approach, n = 3632 samples; cck-escape, n = 1128 samples). **(D-E)** Comparisons of mean dF/F during approaches and escapes from toy rat from samples in the left or right side of the enclosure (the toy rat is located in the right side; left side, n = 5 bins; right side, n = 5 bins; unpaired t-test, *p < 0.05). **(F-G)** Mean dF/F (z-scored) 5-sec before and after approaches, escapes, freeze bouts, and stretch-attend postures in syn-GCaMP6f (F) and cck-GCaMP6f (G) populations. **(H-I)** Mean dF/F (z-scored) in the third of the box near the left wall and the toy rat zone (two-thirds of the environment containing the toy rat) in syn-GCaMP6 (H) and cck-GCaMP6 (I) mice (syn, n = 5; cck, n = 6; paired t-test). Mean ± SEM.

## Methods

### Materials Availability

This study did not generate new unique reagents.

### Experimental Model and Subject Details

All procedures conformed to guidelines established by the National Institutes of Health and have been approved by the University of California, Los Angeles Institutional Animal Care and Use Committee.

### Mice

Cck-IRES-Cre Mice (Jackson Laboratory Stock No. 012706) and wild-type C57BL/6J mice (Jackson Laboratory Stock No. 000664) were used for all experiments. Male and female mice between 2 and 6 months of age were used in all experiments. Mice were maintained on a 12-hour reverse light-dark cycle with food and water *ad libitum*. Sample sizes were chosen based on previous behavioral optogenetic studies on defensive behaviors, which typically use 6-15 mice per group. All mice were handled by experimenters for a minimum of 5 days prior to any behavioral task.

### Rats

Male Long-Evans rats (250-400 g) were obtained from Charles River Laboratories and were individually housed on a standard 12-hour reverse light-dark cycle with food and water *ad libitum*. Rats were only used as a predatory stimulus presentation. Rats were handled for several weeks prior to being used and were screened for low aggression to avoid attacks on mice. No attacks on mice were observed in this experiment.

## Method Details

### Viral vectors

#### Optogeneticse

The following adeno-associated viral vectors (AAV) were used in optogenetic experiments and were purchased from Addgene (Watertown, MA):

AAV9-Ef1a-DIO EYFP (Addgene, 27056-AAV9)

AAV9-EF1a-double floxed-hChR2(H134R)-EYFP-WPRE-HGHpA (Addgene, 20298-AAV9)

AAV2.CMV.HI.eGFP-Cre.WPRE.SV40 (Addgene, 105545-AAV2)

AAV9-hSyn-hChR2(H134R)-EYFP (Addgene, 26973-AAV9)

AAV9-hSyn-EGFP (Addgene, 50465-AAV9)

AAV9-FLEX-Arch-GFP (Addgene, 22222-AAV9)

#### Chemogenetics

The following AAVs, used in chemogenetic experiments, were purchased from Addgene:

AAV8-hSyn-DIO-hM4D(Gi)-mCherry (Addgene, 44362-AAV8)

AAV8-hSyn-DIO-mCherry (Addgene, 50459-AAV8)

#### Fiber Photometry

The following AAVs, used in fiber photometry experiments, were purchased from Addgene:

AAV9.Syn.Flex.GCaMP6f.WPRE.SV40 (Addgene, 100833-AAV9)

AAV9.Syn.GCaMP6f.WPRE.SV40 (Addgene, 100837-AAV9)

### Surgeries

Ten-week-old mice were anaesthetized with 1.5-3.0% isoflurane and affixed to a stereotaxic apparatus (Kopf Instruments). A scalpel was used to open an incision along the midline to expose the skull. After performing a craniotomy, 40 nL of virus was injected into the lateral and ventrolateral (l/vlPA)G (unilateral and counterbalanced for optogenetic activation and fiber photometry experiments, bilateral for inhibition experiments) using a 10 uL Nanofil syringe (World Precision Instruments) at 40 nL/min. Affixed to the syringe is a 33-gauge beveled needle, and the bevel was placed to face medially. The syringe was slowly retracted 11 minutes after the start of the infusion. For l/vlPAG, infusion location measured as anterior-posterior, medial-lateral, and dorso-ventral from bregma were -4.92mm, ± 1.25mm, -2.88mm using a 15-degree angle. For dorsolateral periaqueductal gray (dlPAG), -4.75mm, -0.45mm, -1.9mm using no angle. For chemogenetic experiments, mice received 40nL of AAV8-hSyn-DIO-hM4D(Gi)-mCherry or AAV8-hSyn-DIO-mCherry bilaterally. For optogenetic activation of cck-l/vlPAG cells, 40nL of AAV9-Ef1a-DIO-EYFP or AAV9-EF1a-DIO-ChR2(H134R)-EYFP-WPRE-HGHpA was delivered unilaterally to the l/vlPAG (counterbalancing left or right l/vlPAG) of cck-cre mice. For optogenetic activation of synapsin-expressing l/vlPAG neurons, 40nL of a viral cocktail (1:1) containing AAV2.CMV.HI.eGFP-Cre.WPRE.SV40 and AAV9-EF1a-DIO-hChR2(H134R)-EYFP-WPRE-HGHpA or a viral cocktail containing AAV2.CMV.HI.eGFP-Cre.WPRE.SV40 and AAV9-Ef1a-DIO EYFP was delivered to the l/vlPAG of wild-type mice. For optogenetic activation of synapsin-expressing dlPAG neurons, 40 nL of AAV9-hSyn-EGFP or AAV9-hSyn-ChR2(H134R)-EYFP was delivered unilaterally to the dlPAG of wild-type mice. For optogenetic inhibition of cck-l/vlPAG cells, 40nL of AAV9-Ef1a-DIO EYFP or AAV9-FLEX-Arch-GFP was delivered bilaterally to l/vlPAG. Mice used in optogenetic experiments received a fiber optic cannula (0.22 NA, 200 mm diameter; Doric Lenses) 0.2mm above viral infusion sites. For photometry recordings of cck-l/vlPAG cells, 40nL of AAV9.Syn.Flex.GCaMP6f.WPRE.SV40 was injected into the l/vlPAG of cck-cre mice and an optical fiber was implanted (0.48 NA, 400 mm diameter; Neurophotometrics) 0.2mm above the injection site. For recordings of synapsin-expressing l/vlPAG cells, the same procedure was repeated using AAV9.Syn.GCaMP6f.WPRE.SV40 in wild-type mice. Dental cement (The Bosworth Company, Skokie, IL, USA) was used to securely attach the fiber optic cannula to the skull. Half the mice in each cage were randomly assigned to YFP/mCherry control or ChR2/Arch/hM4di groups. Only mice with opsin expression restricted to the intended targets were used for behavioral assays.

### Immunostaining for NeuN

Fixed brains were kept in 30% sucrose at 4°C overnight, and then sectioned on a cryostat (40 um slices). Sections were washed in PBS and incubated in a blocking solution (3% normal donkey serum and 0.3% Triton-X in PBS) for 1 hour at room temperature. Sections were then incubated at 4°C for 16 hours with polyclonal anti-NeuN antibody made in rabbit (1:500 dilution) (CAT# NBP1-77686SS, Novusbio) in blocking solution. Following primary antibody incubation, sections were washed in PBS 3 times for 10 minutes per wash, and then incubated with anti-rabbit IgG (H+L) antibody (1:1000 dilution) conjugated to Alexa Fluor 594 (red) (CAT# 8889S, cellsignal.com) for 2 hours at room temperature. Sections were washed in PBS 3 times for 10 minutes per wash, incubated with DAPI (1:50,000 dilution in PBS), washed again in PBS and mounted in glass slides using PVA-DABCO (Sigma).

### vGlut2 Immunohistochemistry and histology

Fixed brains were kept in 30% sucrose at 4°C overnight, and then sectioned on a cryostat (40 um slices). Sections were washed in PBS-T (0.3% Triton-X) and incubated in a blocking solution (5% normal donkey serum and 0.3% Triton-X in PBS) for 1 hour at room temperature. Sections were then incubated at 4°C for 16 hours with polyclonal anti-VGLUT2 antibody (#AGC-036, Alomone Labs) made in rabbit (1:500 dilution) in blocking solution. Following primary antibody incubation, sections were washed in PBS-T 3 times for 10 minutes per wash, and then incubated with anti-rabbit IgG (H+L) antibody (1:1000 dilution) conjugated to Alexa Fluor 594 (red) (CAT# 8889S, cellsignal.com) in blocking solution for 2 hours at room temperature. Sections were washed in PBS-T 3 times for 10 minutes per wash, incubated with DAPI (1:50,000 dilution in PBS), and washed again in PBS-T and mounted in glass slides using PVA-DABCO (Sigma).

### Behavior video capture

All behavior videos were captured at 30 frames/sec in standard definition (640×480) using a Logitech HD C310 webcam. To capture fiber-photometry synchronized videos, both the calcium signal and behavior were recorded by the same computer using custom MATLAB scripts that also collected timestamp values for each calcium sample/behavioral frame. These timestamps were used to precisely align neural activity and behavior.

### Behavioral timeline

The order of behavior assays for optogenetic and fiber photometry experiments is as follows (if applicable): open field, elevated plus maze, latency to enter, real time place preference, toy rat exposure, real rat exposure, and pupil dilation. The order of behavior assays for chemogenetic experiments is as follows: toy rat exposure, live rat exposure, hot plate, cued fear conditioning.

### Light delivery for optogenetics

For all ChR2 experiments, blue light was generated by a 473 nm laser (Dragon Lasers, Changchun Jilin, China) at 3-3.5mW with the exception of pupil dilation recordings in which a 1.5mW power was used to avoid overt escape responses during head fixation. For the Arch experiment, green light was generated by a 532 nm laser (Dragon Lasers), and bilaterally delivered to mice at 5-6.6 mW. A Master-8 pulse generator (A.M.P.I., Jerusalem, Israel) was used to drive the blue laser at 20 Hz. This stimulation pattern was used for all ChR2 experiments. The laser output was delivered to the animal via an optical fiber (200 mmcore, 0.22 numerical aperture, Doric Lenses, Canada) coupled to the fiberoptic implanted on the animals through a zirconia sleeve.

### Open field assay with optogenetics

The open field is a square arena (34× 34 × 34 cm) illuminated to 105 lux. Mice had no prior experience in the arena prior to exposure. Exposures are 8 minutes total, with alternating 2-min epochs of laser off or on (OFF, ON, OFF, ON).

### Latency to enter assay with optogenetics

The latency to enter assay is carried out across two consecutive days in a square (47 ×47 × 36 cm) arena. The arena is illuminated to 80 lux and contains a dark burrow (7 × 13 × 11 cm, 2 lux) in one corner. On Day 1, mice are habituated to the entire arena for 10 min. Only mice that spent more time in the burrow compared to the other three corners continued to Day 2. On Day 2, a transparent barrier is placed in the corner opposite of the burrow to create a Holding Zone (HZ). Mice are placed in the arena for 1 min as a context reminder prior to placement in the HZ. Then, 10 trials are carried out, with five laser-OFF and five laser-ON trials interleaved. Prior to all trials, mice are confined to the HZ for 15 sec prior to barrier removal. For light-ON trials, laser is delivered for the latter five of the 15 seconds and continues until the end of the trial. The start of the trial is marked by barrier removal and ends when the mouse enters the burrow or when 60 sec have passed. If mice enter the burrow, they are given 10 sec in the burrow prior to being returned to the HZ. If mice do not enter the burrow, they are immediately returned to the HZ. The procedure is similar between ChR2 activation and Arch inhibition in the LTE, except for laser wavelength and pulse length.

### Place aversion test with optogenetics

Mice were placed in a two-chamber context (20 × 42 ×27 cm) for 10 minutes to freely explore the environment. Both chambers are identical. The following day, mice were introduced into the two-chamber context and blue laser was delivered to the l/vlPAG of cck-cre mice expressing either ChR2 or YFP (20 Hz, 5ms pulses) when they occupied one of the chambers. Laser stimulation was only delivered during exploration of the stimulation chamber. The amount of time mice explored both chambers was tracked across both the baseline and stimulation epochs.

### Elevated plus maze with optogenetics and fiber photometry

The arms of the elevated plus maze (EPM) were 30 × 7 cm. The height of the closed arm walls were 20 cm high. The maze is elevated 65 cm from the floor and is placed in the center of the behavior room away from other stimuli. Arms are illuminated to 8-12 lux. Mice are placed in the center of EPM facing a closed arm. For optogenetic experiments, blue light was delivered in alternating 2-min epochs for five epochs, totalling 10-min exposure. The fifth epoch was excluded from analyses. ChR2 activation was only run in l/vlPAG-cck group, as overt freezing in l/vlPAG-syn group would not be informative and the assay is not feasible with robust jumping exhibited by ChR2 activation of dlPAG-syn mice. For fiber photometry recordings, mice were free to explore the EPM for 10-min.

### Pupil size measurements with optogenetics

Pupil size was measured with the same set-up and methods described previously (Lovett-Barron et al., 2017) Briefly, a camera (AVT Manta, G-032B) coupled to a24 mm/F1.4 lens was used to image the eye under infrared illumination (Thorlabs M780F2). Video was acquired at 60-Hz using pymba, a Python wrapper for AVT camera control. Frame acquisition times and the behavioral task were synchronized with a National Instruments DAQ (NIPCIe-6323). Pupil size was measured from the video using custom-written MATLAB scripts. Each trial lasted 30 s. A 473-nm laser (1.5mW) was delivered to the l/vlPAG of cck-cre mice at 20 Hz, 5ms pulses for 10 s following a 10 s baseline recording. Another 10 s were recorded post-stimulation.

### Live rat exposure assay with optogenetics, chemogenetics and fiber photometry

We used a long rectangular chamber (70 × 25 × 30 cm). Mice were acclimated to this environment for at least two days for 10 minutes each day. During rat exposure, a live rat is restrained to one end of the chamber using a harness attached to a cable with one end taped to the chamber wall. As a behavioral control, we exposed mice to a toy rat (similar in shape and size to a live rat) to assess behavior elicited by visually similar stimuli without actual predatory threat. For the optogenetic experiment, mice were presented with the toy rat for one trial prior to exposure to live rat one day after. On each day, 473-nm laser alternated off or on in 2-min epochs for five epochs, totalling in a 10-min trial. Only the first four epochs (OFF, ON, OFF, ON) were included in analyses. For the chemogenetic experiment, mice were exposed to two toy rat trials on consecutive days followed by two live rat exposures on consecutive days. Mice either received CNO or saline prior to exposures. All trials were 10 mins. For fiber photometry recordings, all mice underwent toy rat exposure followed by live rat exposure the following day for 10 mins each.

### Chemogenetics

Mice used for all chemogenetic experiments (with the exception of cued fear conditioning) were exposed to each threat and control stimuli twice, once following treatment with saline and once following treatment with CNO (10 mg/kg, injected intraperitoneally) 40 minutes prior to the experiment. Only one control or threat-exposure assay was performed per day with each mouse. For cued fear conditioning, all mice received CNO prior to training.

### Heated plate assay with chemogenetics

Heated plate assay was performed on top of a metallic heating plate (14 × 14 cm) (Faithful Magnetic Stirrer model SH-3) across two sequential days on mice expressing AAV8-hSyn-DIO-mCherry or AAV8-hSyn-DIO-hM4D(Gi)-mCherry. On both days, mice received either CNO (i.p., 10 mg/kg) or saline 40 min prior to heated plate exposure. The order of drugs was counterbalanced. Plate is heated to 50 degrees Celsius and enclosed by tall, transparent walls (14×14×24 cm). Mice were closely monitored for pain response. The latency to display a pain-related reaction (hind paw lick or jump) was recorded. All mice showed pain responses within 30 s. After pain response, mice were immediately removed from assay. Pain latency was measured as CNO minus SAL (CNO-SAL).

### Fear conditioning with chemogenetics

Cued fear conditioning was performed across two sequential days on mice expressing AAV8-hSyn-DIO-mCherry or AAV8-hSyn-DIO-hM4D(Gi)-mCherry. On Day 1 (Training), all mice received CNO (i.p., 10mg/kg). Forty minutes later, mice were exposed to a context consisting of metal bar flooring and bare grey walls, and cleaned with 70% ethanol. Training context was illuminated with warm-colored lighting at 40-lux. Mice received ten CS-US pairings pseudo-randomly presented across a 14-min trial. CS was a 10-s pure tone at 70dB and US was a 1-s, 0.6mA shock. CS and US coterminated. The first CS-US pairing began 100s after the start of the trial. On Day 2 (Retrieval Test), mice were exposed to a context consisting of rounded white walls and grey smooth flooring and cleaned with Strike-Bac (Chino, CA, USA). Retrieval test context was illuminated with blue lighting at 40-lux. Mice received 10 CS-only presentations (same CS configuration as training) across a 14-min trial. Freezing was scored using FreezeFrame 5 (Actimetrics, IL, USA). Freeze bouts were at minimum 0.5-s in duration.

### Fiber photometry

Photometry was performed as described in detail previously (Kim et al., 2016). Briefly, we used a 405-nm LED and a 470-nm LED (Thorlabs, M405F1 and M470F1) for the Ca^2+^-dependent and Ca^2+^independent isosbestic control measurements. The two LEDs were band-pass filtered (Thorlabs, FB410-10 and FB470-10) and then combined with a 425-nm long-pass dichroic mirror (Thorlabs, DMLP425R) and coupled into the microscope using a 495-nm long-pass dichroic mirror (Semrock, FF495-Di02-25 3 36). Mice were connected with a branched patch cord (400 mm, Doric Lenses, Quebec, Canada) using a zirconia sleeve to the optical system. The signal was captured at 20 Hz (alternating 405-nm LED and 470-nm LED). To correct for signal artifacts of a non-biological origin (i.e., photo-bleaching and movement artifacts), custom MATLAB scripts leveraged the reference signal (405-nm), unaffected by calcium saturation, to isolate and remove these effects from the calcium signal (470-nm).

### Perfusion and histological verification

Mice were anesthetized with Fatal-Plus (i.p., Vortech Pharmaceuticals, Dearborn, Michigan, USA) and transcardially perfused with PBS followed by 4% paraformaldehyde. Extracted brains were stored for 12 hrs at 4°C in 4% paraformaldehyde before transfer to 30% sucrose for a minimum of 24 hrs. Brains were sectioned into 40um coronal slices in a cryostat, washed in PBS and mounted on glass slides using PVA-DABCO. Images were acquired using a Keyence BZ-X microscope (Keyence Corporation of America, Itasca, IL, USA) with a 4x, 10x, or 20x air objective.

### Behavioral quantification

To extract the pose of freely-behaving mice in the described assays, we implemented DeepLabCut (Nath et al., 2019), an open-source convolutional neural network-based toolbox, to identify mouse nose, ear and tail base xy-coordinates in each recorded video frame. These coordinates were then used to calculate velocity and position at each time point, as well as classify behaviors such as threat approaches, escape runs, stretch-attend postures, and freeze bouts in an automated manner using custom MATLAB scripts. Freezing was defined as epochs when head and tailbase velocities fell below 0.25 cm/s for a period of 0.33 s. ‘Stretch-attend postures’ were defined as epochs for which (1) the distance between mouse nose and tail base exceeded a distance of approximately 1.2 mouse body lengths and (2) mouse tail base speed fell below 1 cm/s. Approach and escape were defined as epochs when the mouse moved, respectively, toward or away from the rat at a velocity exceeding a minimum threshold of 3 cm/s. All behaviors were manually checked by the experimenters for error.

### Statistics

Unpaired t-tests of ON minus OFF or CNO minus SAL transformations were used unless otherwise stated. Two-tailed t-tests were used throughout with α = 0.05. Correlations were calculated using Pearson’s method. Asterisks in the Figures indicate the p values for the post hoc test. Standard error of the mean was plotted in each Figure as an estimate of variation.

## Data and code availability

Custom analysis scripts are available at https://github.com/schuettepeter/l-vlPAG_ActiveAvoidance. Data is available at https://datadryad.org/stash/share/8pOqcca_5ahGWAZ6APF-67IOf8qYJxgOvpHdD6u_XdE

